# Epigenetic characterization of housekeeping core promoters and their importance in tumor suppression

**DOI:** 10.1101/2023.07.26.550759

**Authors:** Martin Loza, Alexis Vandenbon, Kenta Nakai

## Abstract

There has been extensive research on describing cell type-specific (CTS) regulatory interactions, especially between enhancers and promoters. However, constitutively active interactions between CREs have been less studied. In this research, we elucidate the presence of around 11,000 housekeeping CREs (HK-CREs) and describe their main characteristics. Most of the HK-CREs are located in promoter regions, but contrary to expectations, they are not only the promoters of housekeeping genes and are involved in a broader role beyond housekeeping gene regulation. HK-CREs are conserved regions rich in unmethylated CpG sites. Their distribution across chromosomes highly correlates with that of protein-coding genes, and they interact with a large number of target genes in long-distance interactions. In the context of cancer, we observed a reduction in the activity of a subset of HK-CREs, particularly those located at the end of chromosome 19 and associated with zinc finger genes. We investigated the effect of these genes on samples from diverse cancer subtypes, observing a significant reduction in their expression due to aberrant methylation of their core promoters. Finally, an analysis of more than 5,000 patients from 17 cancer subtypes showed an increase in the survival probability of patients with higher expression of these genes, suggesting them as housekeeping tumor suppressor genes. Overall, our work unravels the presence of ubiquitously active CREs indispensable for the maintenance and stability of cells.

## Introduction

Characterizing the different components within the complex transcriptional regulatory mechanism has become a long-standing topic of research [1, 2]. Thanks to the advances in sequencing techniques and experimental protocols, it is possible to investigate different cis-regulatory elements (CREs), e.g., enhancers and promoters, and their differences and similitudes in numerous cell types with unprecedented resolution [3, 4]. These advances have also made clear that classical discrete definitions of CREs do not necessarily fit the intricate interplay between them. For example, several studies have described CREs that may have both enhancer and promoter functions with similar epigenetic features to either classical marks of enhancers or promoters [2, 5]. In this way, promoters with enhancer capabilities could drive the activation of neighboring genes through promoter-promoter interactions [5-7]. In a recent study, contrary to the classical enhancer-promoter interactions on the regulation of cell type-specific genes, it was demonstrated that housekeeping genes (HKG) are mainly regulated through promoter-promoter interactions [8], delineating the importance of this kind of interplay on the maintenance of cells. These new insights into the regulation of HKG and the intrinsic specificity of the core promoters of HKG to certain gene regulators [9] suggest the existence of ubiquitously active CREs besides the promoters of HKG. However, to our knowledge, the existence of these CREs that are active in all cell types has not been described. In this study, we leveraged the activity by contact (ABC) dataset [10], which provides a list of active CREs and their predicted target genes from 131 cell types and tissues, to investigate ubiquitously active CREs in healthy cell types. We elucidate the existence of such housekeeping CREs (HK-CREs), e.g., CREs active in at least 90% of the healthy cell types, and describe their main features. Through several bioinformatic analyses of multiomics data, we show that HK-CREs resided in highly conserved regions that are rich in unmethylated CpG sites, and, similar to HKG [8], they reside in specific loci within the genome as compared with cell type-specific CREs. In conformity with current knowledge on the regulation of HKG [8, 9], we show that most of these ubiquitously active CREs localize in core promoter regions and interact with neighboring genes in complex long-distance promoter-promoter interactions. However, contrary to expectations, our analysis revealed that the core promoters of HKG, which are inherently assumed to be active in every cell type, were not the only ubiquitously active CREs; the core promoters of HKG only accounted for around 18% of the total number of HK-CREs. We then compared the biological processes of HKG and the rest of the genes with ubiquitously active core promoters revealing an intrinsic association between these sets of genes, e.g., both groups are enriched in similar or identical housekeeping functions, which suggests their complementarity to ensure the proper functioning and maintenance of cells. Finally, we investigated the HK-CREs in diverse cancer cell lines, discovering a reduction in the activity of a subset of these elements. Interestingly, we found that the most affected HK-CREs in cancer cells reside close to the telomere region of chromosome 19 and are inherently related to zinc finger genes. Moreover, we show that while most of these HK-CREs with reduced activity were cancer-specific, there was a set of core promoters found to be inactive in numerous cancer cell lines, suggesting the existence of housekeeping tumor suppressor genes. We leveraged public data from diverse popular cancer projects and found a decrease in the expression of putative housekeeping tumor suppressor genes (ZNF154, ZNF135, ZNF667, and ZNF667-AS1) due to aberrant methylation of their core promoters in several cancer subtypes. Despite potential cancer-specific regulation of these genes during cancer progression, a joint analysis of more than 5,000 patients from 17 cancer subtypes showed a significant increase in the survival probability of patients with high expression of these genes, confirming, to some degree, the existence of housekeeping tumor suppressor genes in the human genome.

Overall, our work unveils key cis-regulatory elements active in multiple healthy cell types that, if lost, could strongly affect the housekeeping biological processes of cells and possibly culminate in cancer development.

## Results

### Section 1. Active housekeeping cis-regulatory elements in the human genome

We investigated the cis-regulatory elements (CREs) from the ABC public dataset [10]. This dataset contains a genome-wide list of active CREs from 131 cell types (Supplementary Table 1), identified from the combination of DNase I hypersensitivity (DNase) or assay for transposase-accessible chromatin sequencing (ATAC-seq) data, with chromatin immunoprecipitation sequencing (ChIP-seq) data for the acetylation of the lysine residue at the N-terminal position 27 of the histone H3 protein (H3K27ac). One asset of this dataset is that it provides a list of predicted interactions between CREs and their surrounding genes within 5 Mbps; interactions are quantified from 3D chromatin conformation (Hi-C) data. After selecting significant CRE-gene interactions and filtering out CREs from cell types or tissues related to cancer (n = 26) or various stimuli (n = 34), we obtained around 8.5 million predicted interactions from around 2 million CREs in 71 healthy cell types (Supplementary Table 1). After merging overlapping regions from different cell types (see *Materials and Methods*), we noticed a set of housekeeping CREs (HK-CREs) active in all cell types, which number was stable provided that a sufficient number of cell types were included (see Supplementary Figure 1 and *Materials and Methods*). Then, we randomly selected 50 healthy cell types for further analysis and left the remaining 21 healthy cell types for validation (see *Section 4* and Supplementary Table 1). We then merged overlapping regions across the selected cell types and filtered out suspicious CREs with epigenetic marks similar to a set of 10,000 randomly selected negative sample (NS) regions; NS regions were defined as regions not overlapping with any CRE from the ABC and the SCREEN projects [4, 10] (see *Materials and Methods*). Finally, we obtained a merged dataset with around 846,000 predicted interactions from 136,365 CREs. Figure 1A shows the distribution of the resulting CREs across the number of cell types they are active in. While most of the CREs are active in a low number of cell types, e.g., cell type-specific (CTS-CREs), we found 10,965 HK-CREs active in at least 90% of the 50 cell types. Interestingly, HK-CREs are predicted to interact with a larger number of target genes over longer distances (median number of targets = 22, median distance = 553,817 bps; Figure 1B) as compared with CTS-CREs (mean number of targets = 1, median distance = 63,411 bps). Surprisingly, the HK-CREs presented a considerably high correlation with protein-coding genes in number (r = 0.979; Supplementary Figure 2) and density (r = 0.882; Figure 1C) as compared with CTS-CREs (r = 0.754 and r = 0.601, respectively; Supplementary Figure 3), implying their importance in the gene regulation process. As expected, because of their presumed housekeeping capacity [11, 12], the loci of HK-CREs are significantly conserved and enriched in unmethylated CpG sites as compared with CTS-CREs and negative sample (NS) regions (Figure 1D and Supplementary Figures 4 and 5). Figure 1E shows an example of the conservations and methylation scores of three HK-CREs across nine tissues (tracks across 30 cell types and tissues can be seen in Supplementary Figure 6), where it can be observed that the loci of the HK-CREs overlap with highly conserved and unmethylated regions within a locus that is rich in CpG sites across samples. In contrast, a randomly selected CTS-CRE from T cells presents different methylation patterns across cell types within a poor CpG locus (Supplementary Figure 6).

**Figure 1.**
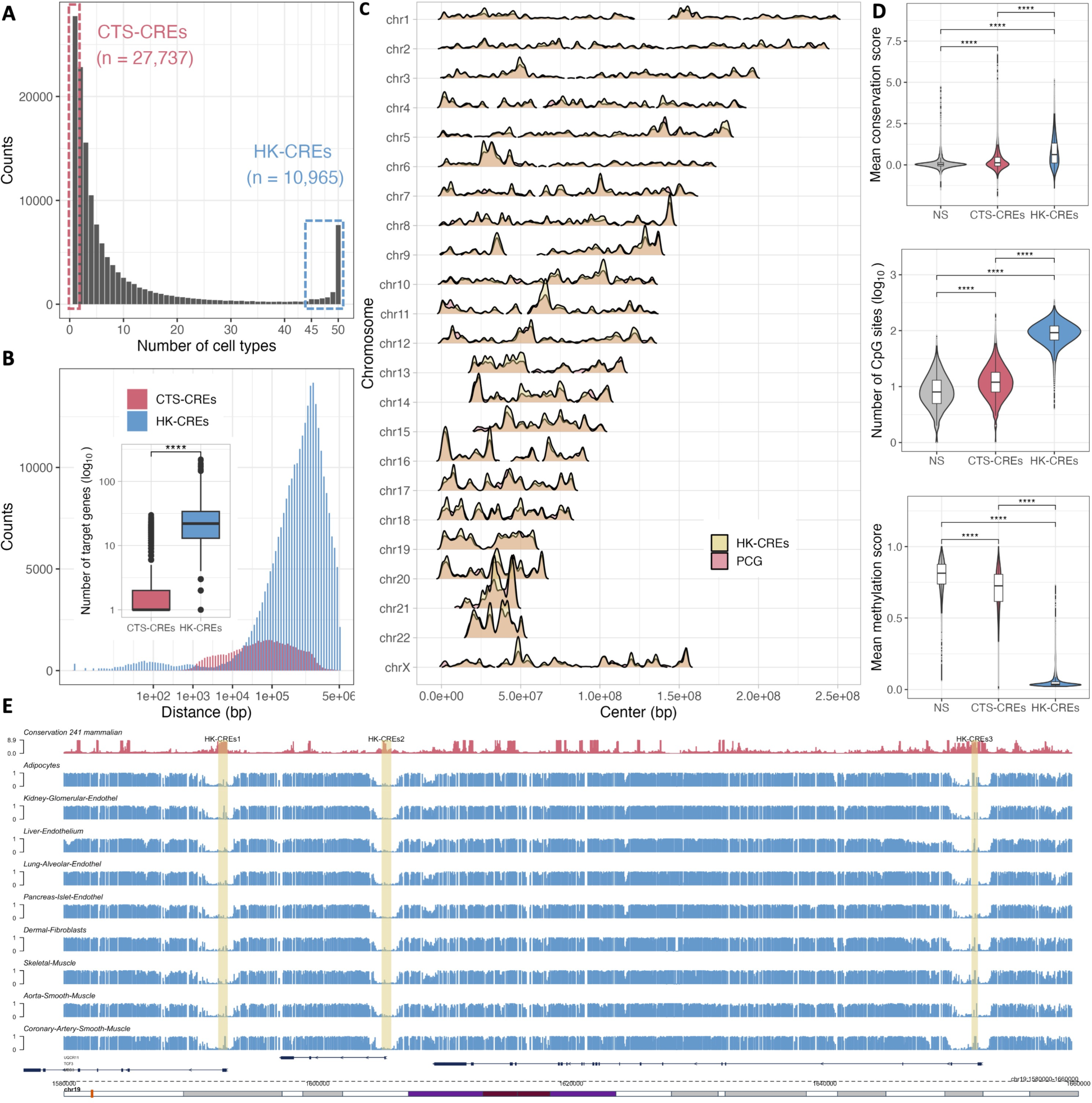
Housekeeping cis-regulatory elements in the human genome. A) After merging overlapping regions from 50 cell types, we observed a significant number of housekeeping cis-regulatory elements (HK-CREs) found to be active in at least 90% of the cells. **B)** HK-CREs interacts with a larger number of surrounding genes in longer distance than CTS-CREs. **C)** The density of HK-CREs highly correlates with the density of protein-coding genes across chromosomes. **D)** HK-CREs reside in conserved regions rich in unmethylated CpG sites. **E)** Conservation scores and methylation levels of three randomly sampled HK-CREs.

### Section 2. Most of the active housekeeping cis-regulatory elements reside in core promoters

We performed an unsupervised classification analysis to further characterize and investigate the functions of housekeeping cis-regulatory elements (HK-CREs). We annotated the CREs with their main epigenetic marks (H3K27ac, H3K4me1, and H3K4me3) and their distance to the nearest transcription start site (dnTSS), using a set of ChIP-seq datasets from the ENCODE project [3](Supplementary Table 2) and the TSS locations from the ENSEMBL project [13], respectively. We used these annotations in an unsupervised classification and visualization process, where we obtained the principal components (PCs) [14] coordinates of CREs and used them as the input data to the Louvain clustering [15] and to the Uniform Manifold Approximation and Projection (UMAP) method [16]. We noticed that the HK-CREs were located at specific parts of the UMAP plot and that they didn’t present a substantial overlap with cell type-specific CREs (CTS-CREs), suggesting their distinctive epigenetic and dnTSS features (Figure 2A-B). The results from clustering confirmed these differences (Figure 2C-D), where CTS-CREs appeared enriched in groups where the number of HK-CREs is scarce and vice versa. Table 1 and Supplementary Figure 7 show the comparison of the feature levels used to manually label nine groups of CREs found by clustering; 5 groups of active CREs: core promoters (CP), promoters (P), enhancer short distance (ESD), enhancers long distance (ELD), and enhancers super long distance (ESLD); one group of inactive enhancers: enhancer short distance inactive (ESDi); two groups of CREs with low signals of epigenetic histone marks: other medium distance (OMD) and other long distance (OLD); and one small group of CREs negative for H3K4me1 (H3K4me1_negative). Interestingly, most HK-CREs were classified as CPs (Figures 2D and 2E) with a “very short” mean dnTSS of 56 bps.

**Figure 2.**
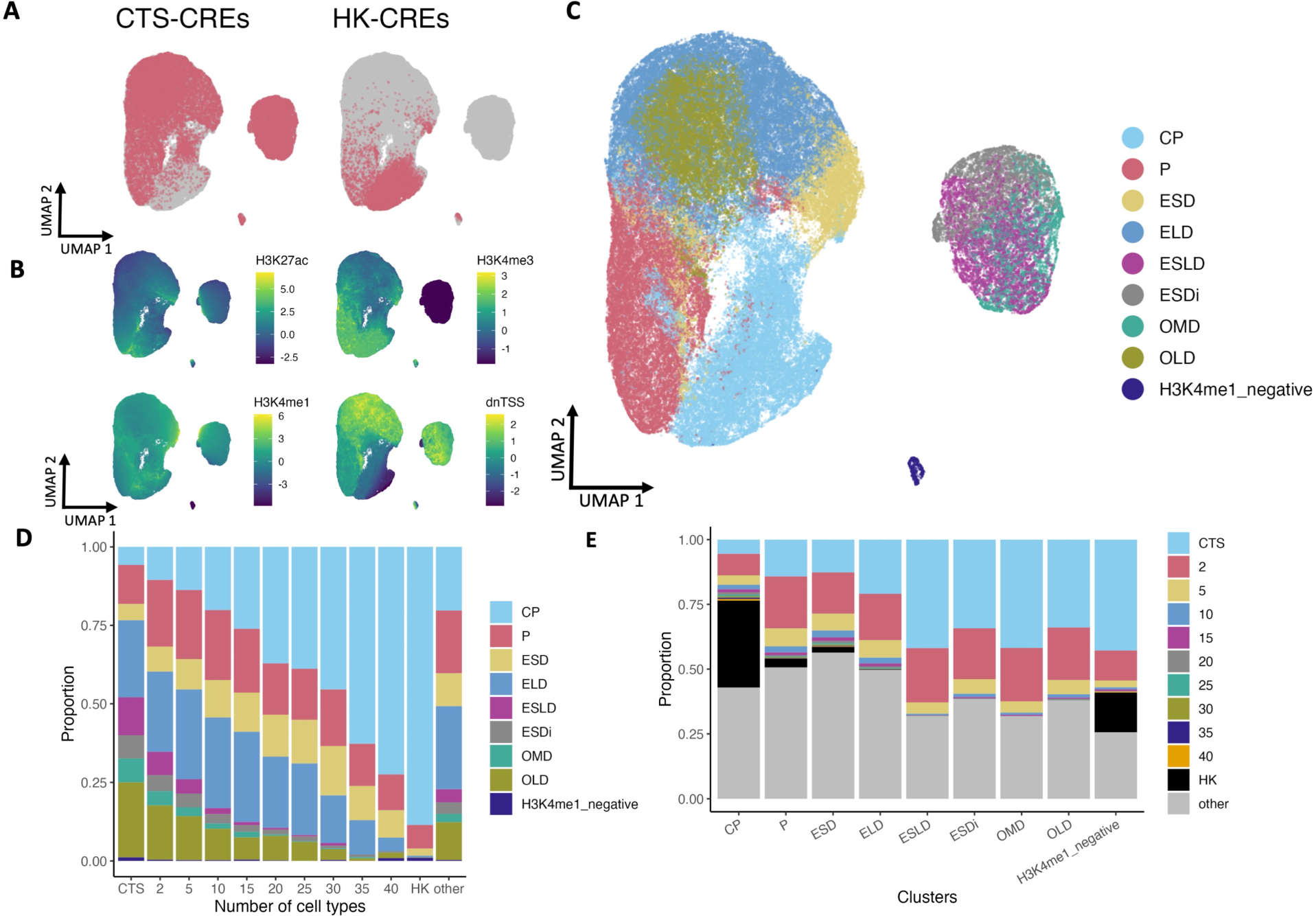
HK-CREs resided in core promoter regions. CREs were annotated with signal levels from H3K27ac, H3K4me3, H3K4me1, and their distance to the nearest TSS (dnTSS). These annotations were used as features in an unsupervised analysis including clustering and visualization. **A-B)** UMAP plot shows a distinct distribution of HK-CREs and cell type-specific-CREs (CTS-CREs) **(A)** with distinct signal levels of histone marks and dnTSS **(B)**. **C-D)** Clustering analysis showed similar results where CTS and HK-CREs were assigned to different clusters. **E)**. Annotation of the clusters using histone signal levels and dnTSS showed that HK-CREs reside primarily in core promoter regions.

**Table 1.** Characterization of clusters. The annotations of clusters were manually assigned using a combination of their epigenetics signals and their distance to the nearest TSS.

| Group | H3K27ac | H3K4me | H3K4me1 | Distance to TSS | Counts | Percentage of total |
| --- | --- | --- | --- | --- | --- | --- |
| **Core promoters (CP)** | + | + | - | Very short | 28,968 | 21.20% |
| **Promoters (P)** | ++ | ++ | - | Short | 24,233 | 17.73% |
| **Enhancers short distance (ESD)** | + | +/- | ++ | Intermediate | 11,445 | 8.37% |
| **Enhancers long distance (ELD)** | +/- | - | + | Long | 32,564 | 23.83% |
| **Enhancers super long distance (ESLD)** | + | -- | + | Very long | 8,063 | 5.90% |
| **Enhancer short distance inactive (ESDi)** | - | -- | + | Intermediate | 5,975 | 4.37% |
| **Other medium distance (OMD)** | - | -- | - | Intermediate | 5,109 | 3.73% |
| **Other long distance (OLD)** | - | -- | -- | Long | 19,563 | 14.31% |
| **H3K4me1 negative (H3K4me1_negative)** | +/- | +/- | Zero | Long | 745 | 0.56% |

### Section 3. Genes whose core promoter is a HK-CRE complement the biological processes of housekeeping genes

Housekeeping genes (HKGs) have been widely investigated as constitutively expressed genes in any cell type regardless of their function or developmental stage [17]. In contrast, to our knowledge, ubiquitously active cis-regulatory elements (CREs) are not clearly defined. We hypothesize that these two elements are closely related, especially for the housekeeping CREs identified as core promoters, e.g., HK-CREs active in more than 90% of the 50 cell types and very close to a transcription start site (TSS). To get an insight into the relationship between our identified housekeeping core promoters (HK-CPs) and housekeeping genes, we obtained the list of genes whose TSS is within 300 bps from the center of an HK-CP and compared them with a set of known 2,131 HKGs [17]. As expected, most of the core promoters of HKGs were included in our set of HK-CPs (Figure 3A). However, our list included the core promoters of a considerable number of other genes (n = 8,681) that haven’t been identified as HKGs, e.g., non-housekeeping genes (nonHKGs) whose core promoter is constitutively active in all cells (an HK-CP) and predicted to interact with surrounding genes by the ABC method. This suggests that, regardless of the activity of their core promoter, the gene expression levels of nonHKG would be different from those of the HKGs. To verify this, we collected RNA sequencing (RNA-seq) data from 37 cell types from the ENCODE project (Supplementary Table 2) and compared the expression levels of HKGs and nonHKGs. Figures 3B and 3C show the mean gene expression of these sets of genes across cell types and the number of cell types expressing them. As expected, the mean expression and the number of cell types expressing nonHKGs were significantly lower than those of HKGs. However, we argued that nonHKGs, whose core promoter is ubiquitously active, could still play an essential role in cell housekeeping processes regardless of their low RNA expression levels. Therefore, we analyzed the biological process (BP) of HKGs and nonHKG from the Gene Ontology project [18]. Interestingly, the analysis showed a high overlap between the BPs of HKGs and nonHKGs with different significance and enrichment levels (Supplementary Figures 8 and 9), suggesting the complementarity of these two sets of ubiquitously active genes. We then performed a differential analysis of the significance and enrichment levels between overlapping BPs of HKGs and nonHKG (max FDR = 0.05). Figures 3D and 3E show the top 10 resulting BPs by significance and enrichment, respectively. Interestingly, the significance analysis (Figure 3D and Supplementary Figure 8) showed that, while HKGs were highly associated with general metabolic, localization, translation, and transport BPs within the cell, nonHKG were related to more specific metabolic and transcription processes, supporting the hypothesized complementarity between these two sets of genes. Surprisingly, the enrichment analysis showed that BPs related to tRNA and regulation of RNA polymerase II, important BPs within the transcription process, were enriched almost exclusively for nonHKGs (Figure 3E and Supplementary Figure 10). Overall, these results confirm the importance of housekeeping core promoters in the gene regulation process and could explain, to some degree, the high correlation between these cis-regulatory elements and the protein-coding genes (Figure 1C and Supplementary Figures 2 and 10).

**Figure 3.**
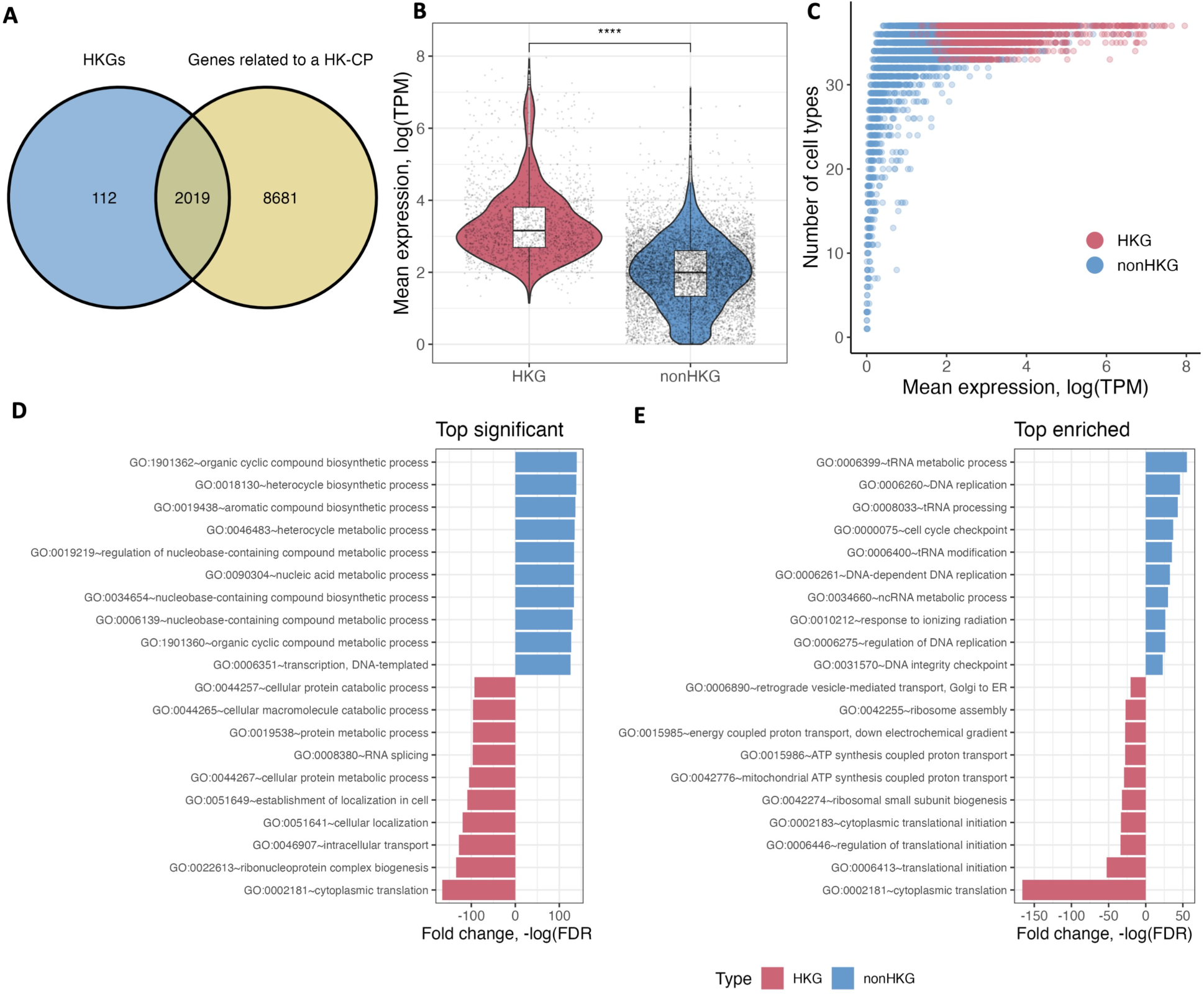
Genes whose core promoter is an HK-CP are significantly enriched in housekeeping biological processes. The genes whose TSS is less than 300 bps from a housekeeping core promoter (HK-CP) were listed and compared with housekeeping genes (HKG). **A)** As expected, most of the core promoters of HKG are HK-CP, however, the core promoters of another 8,681 genes were listed as HK-CPs, e.g. non-housekeeping genes (nonHKG). **B)** Mean gene expression levels of HKGs and nonHKGs over a set of 37 healthy cell types; HKGs present a significantly higher expression than nonHKG. **C)** Mean gene expression levels and the number of cells expressing HKGs and nonHKG; HKGs are ubiquitously expressed in most of the cells, in contrast, nonHKGs present a wider distribution. Besides the differences in gene expression levels and the number of cells expressing HKG and non HKGs, the top biological processes (BPs) sorted by significancy **(D)**, e.g. enrichment of genes in general BPs, and by enrichment (**E)**, e.g. enrichment of genes in specific BPs, suggested the complementarity of these two sets of genes in important housekeeping biological processes.

### Section 4. Inactive housekeeping core promoters in cancer

In our definition of housekeeping cis-regulatory elements (HK-CREs) in *Section 1*, we labeled putative HK regions as those loci active in at least 90% of 50 randomly selected healthy cell types from the ABC dataset. To further validate our list of HK-CREs, we evaluated them using the rest of the 21 healthy cell types and 26 cancer cell lines from the same ABC dataset (Supplementary Table 1). Thus, for each HK-CRE, we calculated the percentage of cells in which it was found to be active in the healthy-validation and cancer datasets. Figure 4A shows the results from this analysis for the set of HK-CREs annotated as core promoters (HK-CPs; similar results for other annotated HK-CREs are shown in Supplementary Figure 11), where the gray dashed line represents our 90% threshold used to define putative housekeeping elements in *Section 1*. Most HK-CPs fulfilled this threshold in healthy cell types (96.94% of HK-CPs, n = 9,411), validating them as housekeeping elements. However, we observed a significant reduction in the distribution of HK-CPs in cancer cell lines (86.05% of HK-CPs, n = 8,354), where 1,354 HK-CPs didn’t fulfill the given housekeeping threshold; Supplementary Table 3 contains the list of HK-CPs that didn’t satisfy the threshold ranked by their ascending percentage of cancer cells they are found active on. To further explore these findings, similar to the analysis performed in *Section 3*, we obtained the list of genes whose TSS is within 300 bps from the center of a HK-CP, and compared the gene expression levels of these genes between a set of 13 cancer cell lines from the ENCODE project [3] and the healthy set used in *Section 3* (n = 37) (Supplementary Table 2). While most of the genes showed a significant correlation between their expression levels in cancer and healthy cells (Figure 4B), the genes whose core promoter was found active in less than 35% of cancer cells (core promoters, n = 19; genes, n = 25) presented a considerably lower median gene expression in cancer cells as compared with healthy cell types (red points in Figure 4B; Supplementary Figure 12). Chromosome 19 was the most affected chromosome with nine HK-CPs active in less than 35% of cancer cells (Supplementary Figure 13 and Table 2). Interestingly, most of the HK-CPs with reduced activity in this chromosome tend to localize towards telomere regions, with the top 3 inactive HK-CPs residing in the region less than 2Mbps away from the end of this chromosome, e.g., near the telomere region, suggesting the importance of this locus for maintaining the integrity of cells.

**Figure 4.**
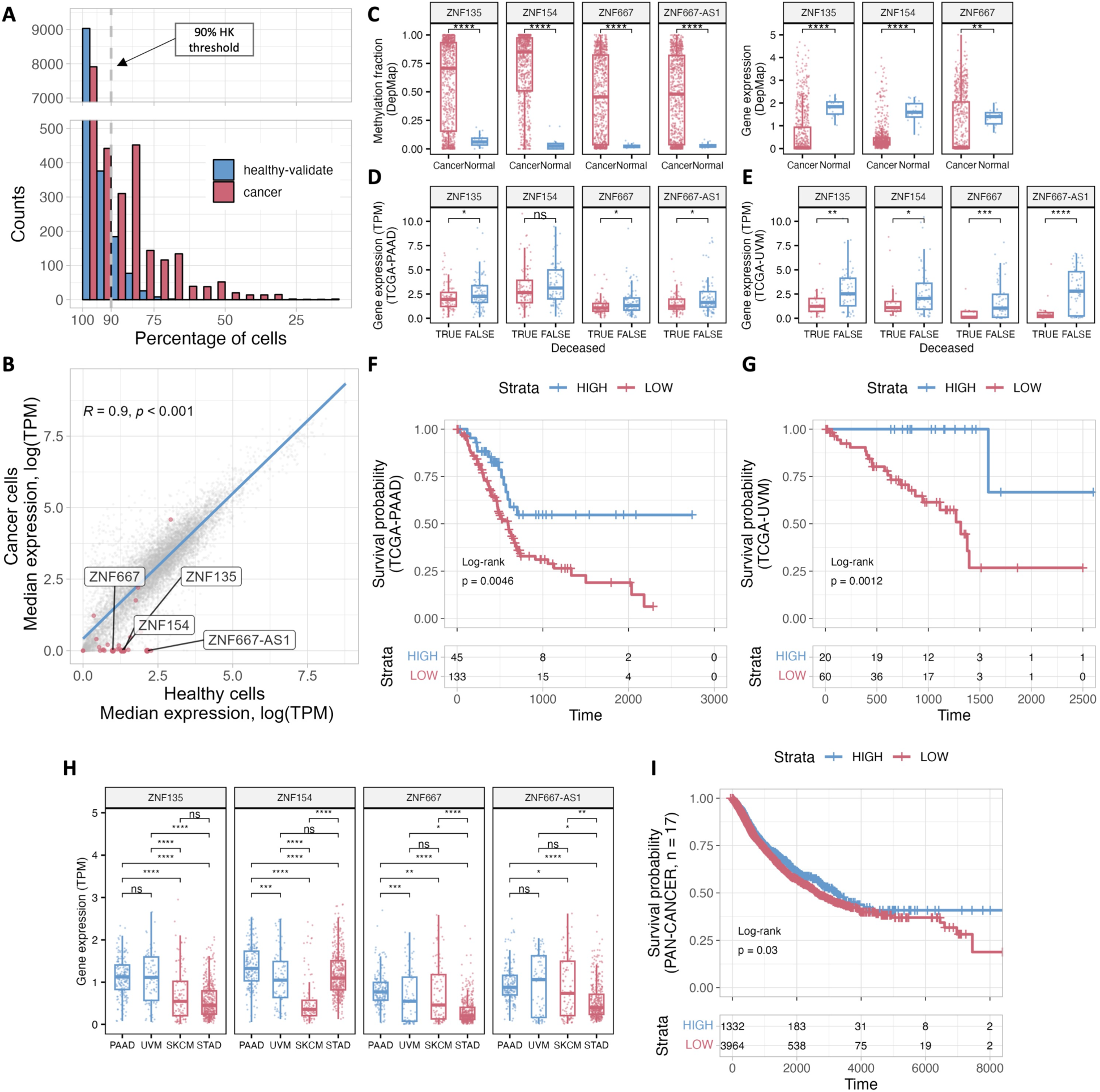
Inactive HK-CPs in cancer cell lines. HK-CP regions were compared with a set of 21 healthy cell types (healthy-validate) and a set of 26 cancer cell lines**. A)** While most of the HK-CPs were found active in more than 90% of the healthy cell types (indicated by the gray dashed line), cancer cells presented a considerable number of inactive HK-CPs; 1,354 HK-CPs were found in less than 90% of the cancer cells. **B)** The expression levels of genes which core promoter is a HK-CP were compared between cancer and healthy samples. In general, there is a highly significant correlation between expression levels in healthy and cancer samples. However, the set of genes which HK-CPs were found active in less than 35% of cancer cells (red-colored dots) presented lower levels of expression on cancer samples. Because of their inactivity in cancer, the genes related to the top 3 inactive HK-CRs are investigated as putative housekeeping tumor suppressor genes: ZNF154, ZNF135, ZNF667, and ZNF667-AS1. **C)** An increased methylation fraction of the promoter regions of putative housekeeping tumor suppressor genes is observed in cancer cells (left), which is related to a decrease in their gene expression levels (right). Further analysis of these genes in cancer samples from TCGA revealed significant changes among deceased conditions in diverse cancer subtypes such as pancreas adenocarcinoma **(D)** and uveal melanoma **(E)**. **(F-G)** Independent survival analyses on these projects revealed a significantly higher survival probability of samples with high mean expression of these genes. **H)** Different patterns of expression levels of samples from distinct projects might suggest cancer specific regulation of these genes. **I)** After normalizing the differences between samples from different projects, a joint survival analysis including more than 5,000 samples from 17 different cancer subtypes is performed. A relatively significant increased survival probability is observed in samples with a high expression of the putative tumor suppressor genes.

**Table 2.** Top inactive HK-CPs in cancer cell lines. Inactive HK-CPs found in less than 35% of cancer cells ranked by their distance to the end of the chromosome they reside in, e.g. telomere regions.

| HK-CP Region | Gene name | TSS | % cancer cell lines | % healthy cell types | Distance to the end of the chromosome |
| --- | --- | --- | --- | --- | --- |
| chr19:58058989-58059488 | **ZNF135** | 58,059,239 | 11.53 | 95.23 | 558,377 |
| chr19:57935138-57935637 | **ZNF418** | 57,935,387 | 30.76 | 100.00 | 682,229 |
| chr19:57708962-57709461 | **ZNF154** | 57,709,204 | 11.53 | 95.23 | 908,412 |
| chr19:57583726-57584309 | **ZIK1** | 57,584,140 | 26.92 | 100.00 | 1,033,476 |
| chr19:56507593-56508092 | **ZNF471** | 56,507,850 | 30.76 | 95.23 | 2,109,766 |
| chr19:56477152-56478177 | **ZNF667-AS1** | 56,477,590 | 15.38 | 95.23 | 2,140,026 |
| chr19:56477152-56478177 | **ZNF667** | 56,477,374 | 15.38 | 95.23 | 2,140,242 |
| chr19:53254646-53255153 | **ZNF677** | 53,254,898 | 23.07 | 95.23 | 5,362,718 |
| chr19:51887687-51888274 | **ZNF649-AS1** | 51,888,025 | 26.92 | 95.23 | 6,729,591 |
| chr19:51887687-51888274 | **ZNF577** | 51,887,980 | 26.92 | 95.23 | 6,729,636 |
| chr19:36605027-36605554 | **ZNF382** | 36,605,313 | 34.61 | 90.47 | 22,012,303 |
| chr19:36605027-36605554 | **ZNF529** | 36,605,276 | 34.61 | 90.47 | 22,012,340 |

### Section 5. Potential housekeeping tumor suppressor genes

We further investigated the genes with the top 3 inactive core promoters: ZNF154, ZNF135, ZNF667-AS1, and ZNF667, using public data from commonly used cancer databases. We performed a pan-cancer analysis, e.g., joint analysis of diverse tumor samples, using the TNMplot web tool [19] for differential gene expression analysis of tumor versus normal samples. We observed similar results to those presented in the previous section (Figure 4B), where the expression levels of these four genes were lower in tumor samples in various cancer subtypes (Supplementary Figures 14 and 15). To confirm the effect on the core promoter regions of these genes in cancer, we compared the methylation fraction around the promoter regions (Figure 4C left) and the expression levels of these genes (Figure 4C right) for diverse cancer subtypes from the DepMap portal [20]; expression levels of ZNF667-AS1 were not available at the time of the analysis. Surprisingly, the promoter regions were significantly more methylated in cancer samples, whereas normal samples presented almost no methylation. The aberrant methylation of these HK-CPs explains, to some degree, their observed inactivity (see *Section 4* and Figure 4A) and the low expression levels of the genes they regulate in cancer samples (Figure 4B, Figure 4C right, and Supplementary Figures 14 and 15).

Because of the ubiquitous activity of their core promoters in healthy cell types and their inactivity in most of the investigated cancer cell lines (due to aberrant methylation), we hypothesized that these four genes: ZNF154, ZNF135, ZNF667-AS1, and ZNF667, could be related to housekeeping tumor suppression functions. Surprisingly, the tumor suppression capacity of these genes has already been observed for different cancer subtypes in diverse tissues: non-small cell lung cancer, renal cancer carcinoma, thyroid cancer, nasopharyngeal carcinoma, squamous cell carcinoma, uveal carcinoma, etc. [21-45]. Moreover, the attenuation of the anti-tumor capacity of these genes has been related to the aberrant methylation of their core promoters in individual [36, 39, 45] and pan-cancer analyses [40, 43, 44].

To further confirm our hypothesis of the existence of housekeeping tumor suppressors, we performed a comprehensive data analysis of 17 cancer projects from The Cancer Genome Atlas Program (TCGA) (Supplementary Table 4). We first compared the gene expression levels of samples, e.g., samples from patients diagnosed with cancer, from different deceased conditions across projects (Supplementary Figure 16). While most of the projects presented non-significant variations, the UVM (eye and adnexa; uveal melanoma) and PAAD (pancreas; pancreatic adenocarcinoma) projects presented significant distinctions (Figures 4D-E and Supplementary Figure 16), where the expression levels of the putative tumor suppressor genes were higher in non-deceased samples, suggesting a higher survival probability of patients from these samples. Then, for each project, we performed a joint survival analysis [46] of the four genes with samples stratified into two groups: strata = HIGH, samples with a mean expression level (see *Methods*) higher than 75% of the samples; strata = LOW, others. As expected, the survival analyses of PAAD and UVM (Figures 4F and 4G, respectively) showed a significantly higher survival probability of HIGH samples. Interestingly, even though the p-values were not significant, we observed HIGH samples to present a higher survival probability in several projects, e.g., BRCA (breast; breast invasive carcinoma), LUAD (bronchus and lung; lung adenocarcinoma), and ACC (adrenal gland; adrenocortical carcinoma; Supplementary Figures 17 and 18). However, some projects, such as the STAD (stomach; stomach adenocarcinoma) and SKCM (skin; skin cutaneous melanoma), presented an inverse effect, where LOW samples showed a higher survival probability (Supplementary Figure 19). Because of the significantly lower expressions of ZNF135, ZNF667, and ZNF667-AS1 genes in samples from stomach cancer as compared with healthy samples (Supplementary Figures 14 and 15), we argue that this inverse effect in STAD and SKCM projects does not entirely contradict our hypothesis of potential housekeeping tumor suppressor genes; instead, it might suggest cancer-specific changes on these genes driven by other latent variables (See Discussion and Supplementary Figure 20). To exemplify this cancer-specific effect, we compared the expression levels of samples from PAAD and UVM (projects that agree with our hypothesis) with samples from SKCM and STAD (projects that disagree with our hypothesis), where for most of the comparisons (Figure. 4H) we observed a significantly lower expression of samples from the latter projects. Given the potential cancer-specific changes of putative housekeeping tumor suppressor genes, we normalized the expression levels of each gene across projects (see *Methods*) to perform a joint survival analysis of the 17 TCGA projects. Figure 4I shows the results of the analysis where HIGH samples (n = 1332), e.g., samples with a mean expression level (see *Methods*) higher than 75% of the samples, presented a significantly higher survival probability as compared with LOW samples (n = 3964; independent analyses for each gene are shown in Supplementary Figure 19). Together, these results suggest the existence of active housekeeping core promoters related to housekeeping tumor-suppressor genes that, when methylated, may culminate in the development of cancer regardless of their malignancy phenotype variant.

## Discussion

Our analysis unveils a key housekeeping element of the gene regulatory mechanism that, to our knowledge, hasn’t been described before. Through several bioinformatics analyses leveraging diverse public epigenetic and gene expression datasets, we reveal the existence of housekeeping cis-regulatory elements (HK-CREs) that are mainly located within core promoter regions and are not necessarily related to housekeeping genes (HKG), e.g., they are not the core promoters or promoters of HKG. We defined HK-CREs by randomly selecting a substantial number (n = 50) of healthy cell types to minimize biases in our analysis, ensuring a representative sample. Moreover, besides using a validation dataset with 21 healthy cell types and a set of 10,000 negative samples to assess our findings, we proved in distinct ways the robust location within the genome of the HK-CREs. For instance, we argue that the probability that most of the overlapping regions from 50 cell types intersect within the loci of core promoters (median distance to TSS = 56bps) is exceptionally low. In addition, our analysis using a methylation atlas from healthy cells [47] showed that HK-CREs are located in dense unmethylated CpG loci for multiple cell types, which can’t just occur by chance. Therefore, we feel confident about the existence and robust location of the HK-CREs.

When we first observed that most of the HK-CREs overlapped with core promoter regions, we first thought that we captured the core promoters of HKG. However, the HKG only represented less than 20% of the genes in our list, e.g., genes which core promoter is a HK-CRE. Our immediate conjecture was that the remaining genes in our list would be genes that have not been correctly annotated as HKG. However, our comparison of gene expression levels showed significant differences between both groups. These results do not contradict our finding of ubiquitously active CREs, as their definition is not based on their transcription levels, but we recognize that there is a bias towards the list of HKG and datasets used in the analysis. Further experimental validations with higher sequencing depth may clarify the distinction between these two sets of genes that seem to complement each other in housekeeping biological processes in the cell.

In this manuscript, we focused on exploring and characterizing HK-CREs instead of the interactions predicted by the ABC method. We observed intricate cooperative interactions between HK-CPs, where various HK-CPs exhibit interactions with one another and with the genes in their proximity. This suggests the presence of cis-regulatory hubs with promoter-promoter interactions in their core forming complex regulation networks. We are aware of the importance of these intriguing promoter-promoter interactions not only for regulating HKGs but also for cell type-specific genes. Nonetheless, these interactions come from predictions and must be verified experimentally, but we believe these two results can be treated separately without conflict.

Regarding transcription factor binding motifs found in HK-CREs, we observed an enrichment of common GC-rich motifs, e.g., SP1 and SP2. As this information somehow overlaps with the conservation and enrichment of CpG sites shown in Figure 1D, we did not explore the enriched motifs further. However, together with appropriate experimental validations, differential analyses of enriched motifs across annotations of HK and CTS-CREs, e.g., core promoters vs. promoter regions, could reveal important motifs for the complex enhancer grammar at different levels.

In our analysis of inactive housekeeping core promoters (HK-CPs) in cancer cell lines (see Section 4), we tried to avoid bias toward the 26 cell lines in the ABC dataset by focusing on the top inactive HK-CPs from a list of 1,354 elements. Moreover, our investigation revealed that the most affected elements group together in the telomere of chromosome 19, which proves, to some degree, the robustness of our results. In addition, for the top 3 inactive HK-CPs in cancer cells, we confirmed their aberrant methylation and the decreased expression levels of the genes they regulate through comprehensive analysis using data from diverse public sources as ENCODE, DepMap, and the web tool TNMplot, obtaining a general agreement among results.

Finally, for the set of putative housekeeping tumor suppressor genes (ZNF135, ZNF154, ZNF667, and ZNF667-AS1), e.g., genes whose HK-CPs are active in less than 20% of the cancer cells, we performed a comprehensive analysis using cancer samples of 17 projects from TCGA. Even though we observed an increased survival probability of samples with high expression levels of these genes in diverse projects, only two were significant: PAAD (pancreas adenocarcinoma) and UVM (uveal melanoma). However, we argue the existence of latent cancer-specific mechanisms that affect these genes which must be considered. For instance, we observed interesting cancer-specific dynamics in the expression levels of these genes across samples from different cancer stages (Supplementary Figure 20). For example, the expression levels from PAAD samples showed a decline from the early to intermediate stages, with a final increased expression in the late stages. These dynamic changes might be related to other biological processes of these genes and the inherent cancer-specific necessity to down/up-regulate them during different stages. Further analysis including samples from the same patient at different stages might reveal the intrinsic mechanism behind these intriguing changes.

Together, even though our results should be verified experimentally, we are confident that they continue to hold significant relevance for pan-cancer studies concerning the characterization of housekeeping cis-regulatory elements involved in tumor suppression.

## Materials and Methods

### Quality control and pre-processing of regions

First, we filtered out not significant regions with an ABC score lower than 0.015 as in the original publication [48]. Then, we lifted up the regions coordinates to the hg38 human genome using the *CrossMap* assembly converter [49] implemented in the Ensemble website [13].

We manually filtered out cancer cell lines or stimulated cell types from the original 131 cells in the ABC dataset [10], obtaining a healthy dataset of 71 cell types and tissues (Supplementary Table 1). To investigate the dependency of the housekeeping (HK) regions on the number of included cell types and on how strictly we define a region as to be HK, we merge the regions (see *Merging process*) with an increasing number of randomly selected cell types using a 70, 80, 90, or 100 % definition of HK, e.g., a given region is an HK region if it is found in at least 70, 80, 90, or 100% of the cell types. We observed that the number and length of the merged HK regions stabilized after we included around 30 to 40 cell types with a 90 or 100% definition (Supplementary Figure 1). Therefore, we randomly selected 50 healthy cell types (Supplementary Table 1) and merged their regions for further analyses. The regions from the remaining 21 healthy cells were kept for validation.

After annotating the main histone marks of the merged regions (see *Annotation of histone marks*), as a quality control step, we filter out regions with histone marks similar to a set of 10,000 negative samples (see *Definition of negative samples*). For this, we clustered regions using the Louvain algorithm [15] implemented in the *igraph* R package [50] using the histone marks and the distance to the nearest TSS as features. Finally, we retained 135,917 regions as putative cis-regulatory elements (see Table 1).

### Merging process

For a given set of cell types, we merged overlapping regions following the next iterative process until no regions were left in the raw dataset:

- <u>Step 1: Merge raw overlapping regions across cell types.</u> We used the function *bedr.merge.region* from the bedr R package [51] with a distance of 20bps.
- <u>Step 2. Select merged regions that overlap on the maximum number of cell types.</u> In the case of housekeeping regions, we merged and selected regions overlapping with a defined 90% percentage of the total number of cells, e.g., regions that overlap in at least 90% of the cell types.
- <u>Step 3. Filter out raw regions overlapping with the selected merged regions.</u> We used the *findOverlaps* function from the GenomicRanges R package [52].
- <u>Step 4. Repeat Step 1 using the filtered raw regions.</u>

### Definition of negative samples

We defined a set of 10,000 negative samples (NSs) as non-overlapping regions with cis-regulatory elements (CREs) from the ABC database (131 cell types) nor the SCREEN project [4] as follows:

- Merge regions from the ABC and SCREEN datasets using the *reduce* function from the GenomicRanges R package [52] with a minimum gap width of 100bps.
- Calculate the gaps between the merged regions using the *gaps* function from the GenomicRanges R package.
- Filter out gaps located in centromeres or at the end of telomeres.
- Filter out gaps less than 3Kbps long. This length intends to keep regions of 1.5Kbps which are at least 500bps away from any extended CRE (see *Annotation of histone marks*).
- Select 10,000 random gaps and define NS as 500bps regions located at the center of the selected gaps.

### Annotation of histone marks

We annotated the H3K27ac, H3K4me1, and H3K4me3 histone marks of an extended version of the cis-regulatory elements and negative samples. The extended version includes the original regions plus 250bps on each flank to account for adjacent histones.

We downloaded ChIP-seq histone marks datasets from the ENCODE project [3] (Supplementary Table 2) of samples selected as being similar to the main 50 healthy cells/tissues (see *Quality control and pre-processing of regions*). Following the pre-processing used on the SCREEN project [4], we performed independent log(signal+1) normalization and scaling of the samples. Finally, we annotated the histone marks of the extended regions as the average signal of overlapping peaks across pre-processed samples; overlapping peaks were selected as those with at least 250bps overlap with the extended regions.

### Annotation of the distance to the nearest TSS

We annotated the distance to the nearest TSS of cis-regulatory elements and negative samples using the ENSEMBL transcripts annotation version GRCh38.107.

### Annotation of the conservation scores

We annotated the mean conservation scores on a fixed-length version of the cis-regulatory elements (CREs). The fixed-length CREs were defined to be 500 bps at both flanking loci from the center of each CRE. Then, we overlapped each fixed-length CRE with the conservation scores of 241 mammals from the Zoonomia project [53] and calculated the mean conservation score as the mean score among overlapping regions.

### Gene Ontology Analysis

Gene ontology analysis of genes was performed using the DAVID web tool [54]. Significant processes were selected to have max FDR <= 0.05. The differential enrichment analysis was performed by calculating the fold change of −log(FDR) between housekeeping and non-housekeeping genes.

### Methylation Atlas

We downloaded the methylation scores from 34 randomly picked samples from the DNA methylation atlas of normal human cell types [47] (Supplementary Table 2). Then, for each fixed-length CRE and NS, we counted the number of overlapping CpG sites and calculated the mean methylation score across cell types; the fixed-length regions were defined as regions of 1kbps concentric with every CRs and NS. To produce the methylation tracks from Figure 1E and Supplementary Figure 6, we used the *track_plot* function from the trackplot R package [55].

### ENCODE RNA-seq datasets

All the healthy and cancer RNA-seq data were obtained from the ENCODE project [3] (Supplementary Table 2). For pre-processing and quality control of samples, we followed the next pipeline:

- Load the TPM data using the *tximport* Bioconductor package [56].
- Filter out genes with zero TPM in all the samples.
- Map the gene labels to match the gene names used in the ABC dataset. We used the *org.Hk.eg.db* and *AnnotationDbi* Bioconductor packages [57, 58] to match the gene names with a combination of ALIAS, ENSEMBL, and ENTREZID.
- Natural log-normalize the TPM data.

### DepMap methylation and RNA-seq data

The methylation and RNA-seq data were downloaded directly from the DepMap website [20]. We used the *secondary_lineage* column in the metadata of the samples and re-labeled “Non-Cancerous” as “Normal” and all the other cells as “Cancer”.

### TCGA clinical and RNA-seq data

TCGA data were downloaded and analyzed using the TCGAbiolinks R package [59]. In this study, we included only projects related to a single primary site (see *Supplementary Table 4*).

For each selected TCGA project (see Supplementary Table 4), the clinical data was obtained by using the *GDCquery_clinic* function and setup as follows:

- Samples were classified as *deceased* = TRUE | FALSE, based on their “Alive” status from the *vital_status* column in the metadata.
- The overall survival of samples was assigned as: if *vital_status* == “Alive” then *overall_survival* = *days_to_last_follow_up*, else *overall_survival* = *days_to_death*, where *days_to_last_follow_up* and *overall_survival* are columns from the clinical metadata.

For each selected TCGA, a query for downloading publicly available RNA-seq data was created using the *GDCquery* function; data.category = “Transcriptome Profiling”, experimental.strategy = “RNA-seq”, workflow.type = “STAR – Counts”, data.type = “Gene Expression Quantification”, sample.type = “Primary Tumor”, access = “open”.

The RNA-seq data was downloaded using their corresponding query and the *GDCdownload* function. The TPM of genes was used in all the analyses.

To calculate the mean expression of each sample used for the survival analyses (Figures 4F-G, Figure 4I, and Supplementary Figures 17-19) we Z-transformed the TPM expression levels of the ZNF154, ZNF135, ZNF667, and ZNF667-AS1 genes on each project. Then, a *mean_expression* was calculated for each sample as the mean value of the transformed expressions.

To perform the survival analyses, we stratified the samples based on their *mean_expression* as follows: if *mean_expression* >= 3^rd^ quantile(*mean_expression* of all samples), then *strata* = “HIGH”, else *strata* = “LOW”. Using the *strata* label, we fit a survival model using the *survfit* function from the survival R package [60]; Surv(*overall_survival*, *deceased*) ∼ *strata*. Finally, we created the survival plots using the *ggsurvplot* function from the survminer R package [61].

### Statistical information

For the statistical comparison used in the manuscript, we used the *stat_compare_means* function from the ggpbur R package [62] with the next convention for symbols indicating statistical significance:

- ns: p > 0.05
- *: p <= 0.05
- **: p <= 0.01
- ***: p <= 0.001
- ****: p <= 0.0001

In Figure 1D and Figure 3B, we used an unpaired Wilcoxon test with a two-sided alternative hypothesis. In Supplementary Figure 12, we used a paired Wilcoxon test with a two-sided alternative hypothesis.

### Other analyses and plots

To plot the density of regulatory regions, protein-coding genes, and transcripts of genes, we used the *geom_density_ridges* function from the ggridges R package [63]; bandwidth = 1e6.

To plot the genomic regions by chromosomes, we used the Chromomap R package [64]; window.size = 1e6.

## Data Availability

The ABC dataset was downloaded from https://www.engreitzlab.org/resources. Histone modification data and RNA data were downloaded from the ENCODE project. The sources and accession numbers are summarized in Supplementary Table 2. Conservation scores were downloaded from the Zoonomia project from https://cglgenomics.ucsc.edu/data/cactus/.

Methylation scores were downloaded from the GEO (accession no. GSE186458). The accession numbers of cell types are summarized in Supplementary Table 2.

Annotations of genes and transcripts were downloaded from the ENSEMBL project http://asia.ensembl.org/Homo_sapiens/Info/Index.

Methylation and RNA-seq data from the DepMap project were downloaded directly from the data explorer of their website; https://depmap.org/portal/interactive/.

TCGA clinical and expression data were downloaded using the TCGAbiolinks R package. The description of the projects is summarized in Supplementary Table 4.

Any additional information generated in this paper is available from the corresponding author upon reasonable request.

## Acknowledgments

The authors thank Prof. Katsuhiko Shirahige and Dr. Takashi Fukaya for their fantastic discussion and invaluable comments.

This work was supported by Grants-in-Aid for Scientific Research (grant number 22K06189 for K. N. and JP22K21301 for M. L.) from the Japan Society for the Promotion of Science.

## Authors’ contributions

M. L. contributed to the conception, design, data acquisition, data analysis and interpretation, drafted and critically revised the manuscript. A. V. and K. N. contributed to the conception, design, interpretation of results, edited and critically revised the manuscript. K. N. supervised the project.

## Competing interests

The authors declared no potential conflicts of interest concerning this article’s research, authorship, and/or publication.

## Notes

### Competing Interest Statement

The authors have declared no competing interest.

### Summary of Updates

Section 4 was expanded and divided into two new sections: Section 4, describes housekeeping core promoters inactive in cancer cell lines; Section 5, explores the top inactive housekeeping core promoters as regulatory elements of putative housekeeping tumor suppressor genes.

