## Supplementary Figures for "Epigenetic characterization of housekeeping core promoters and their importance in tumor suppression"

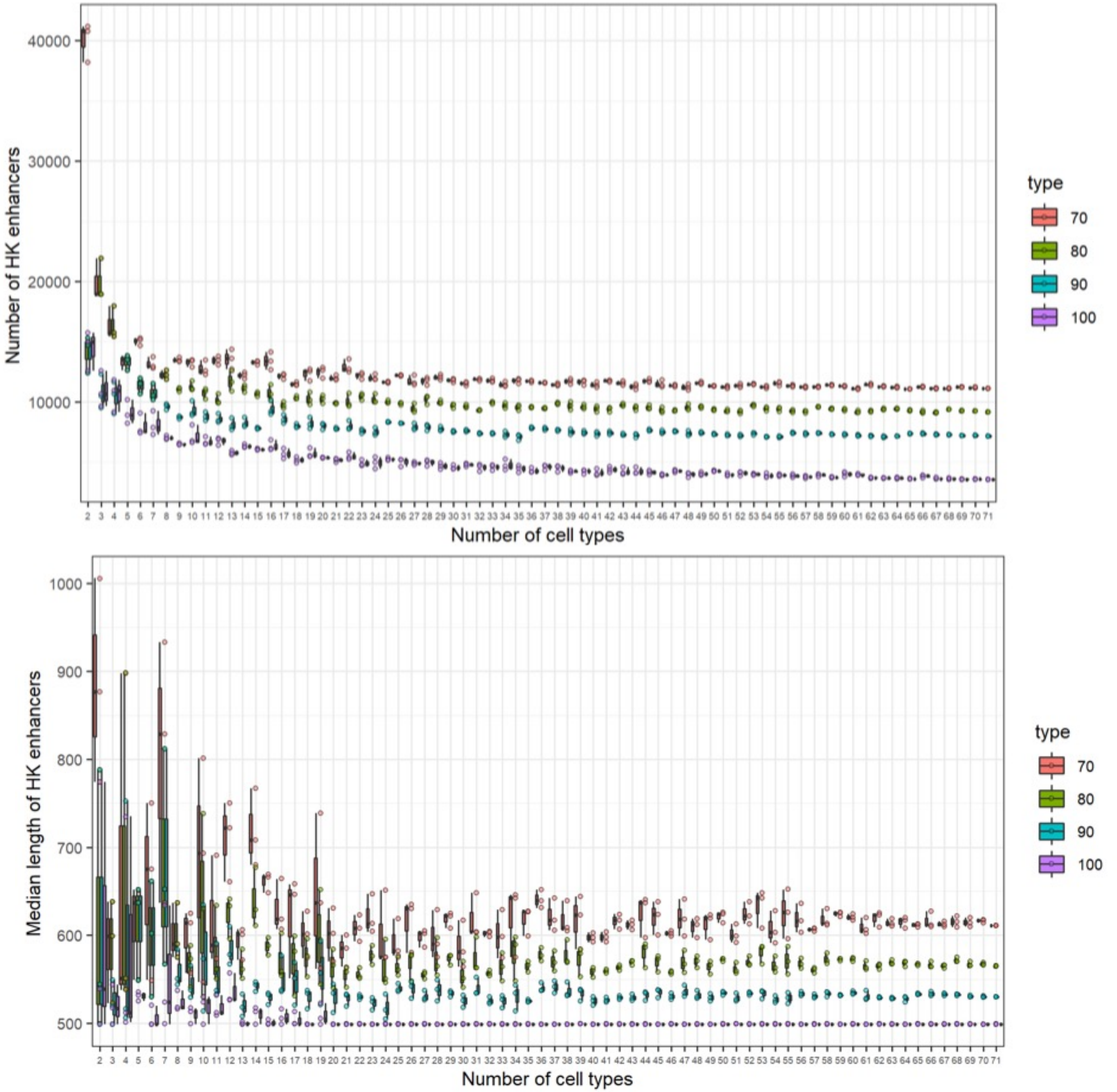


**Supplementary Figure 1.** The dependency of the HK-CREs on the number of selected healthy cell types and on the definition of housekeeping elements was investigated, e.g., a CRE is considered a housekeeping CRE if found in at least 90% of the cells. In general, the distribution of the number of HK-CREs and their length stabilize after including around 30-40 cell types using a 90% definition threshold.

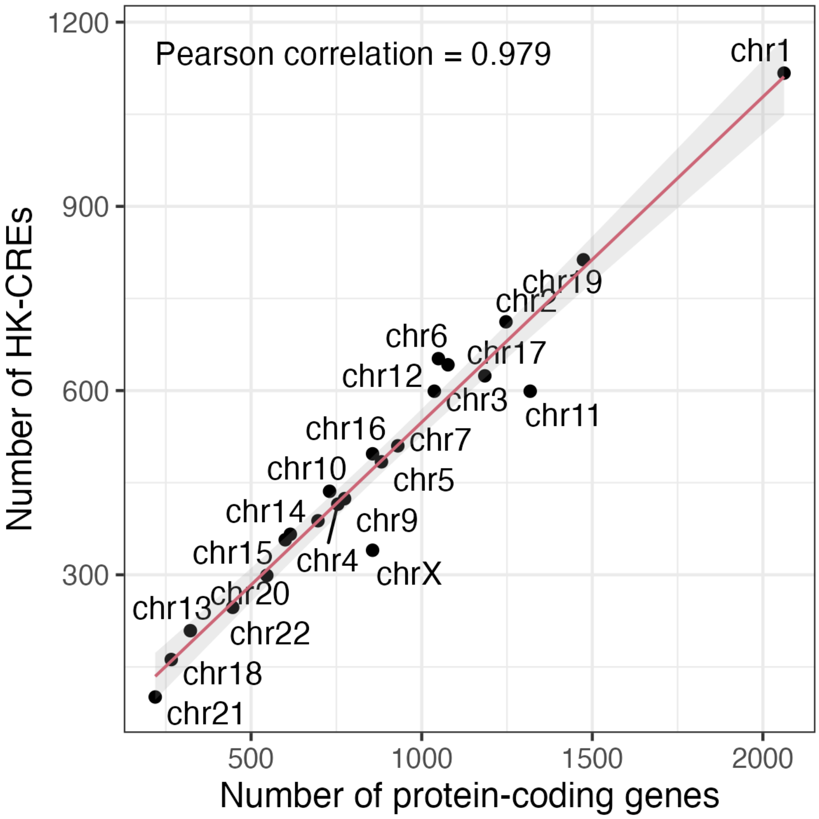


**Supplementary Figure 2.** The number of HK-CREs highly correlate with the number of protein coding genes across chromosomes.

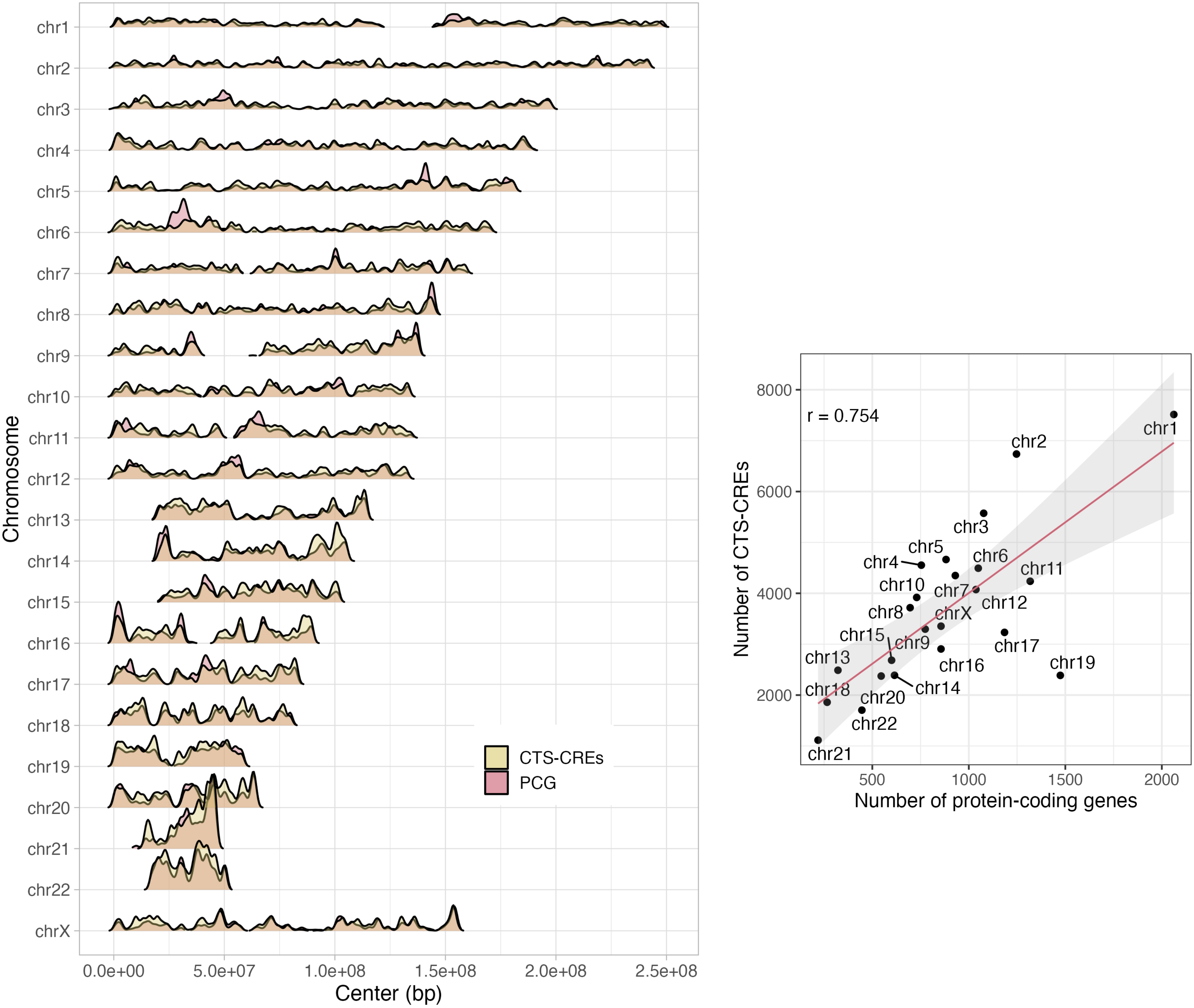


**Supplementary Figure 3.** Number and density of CTS-CREs as compared with protein-coding genes.

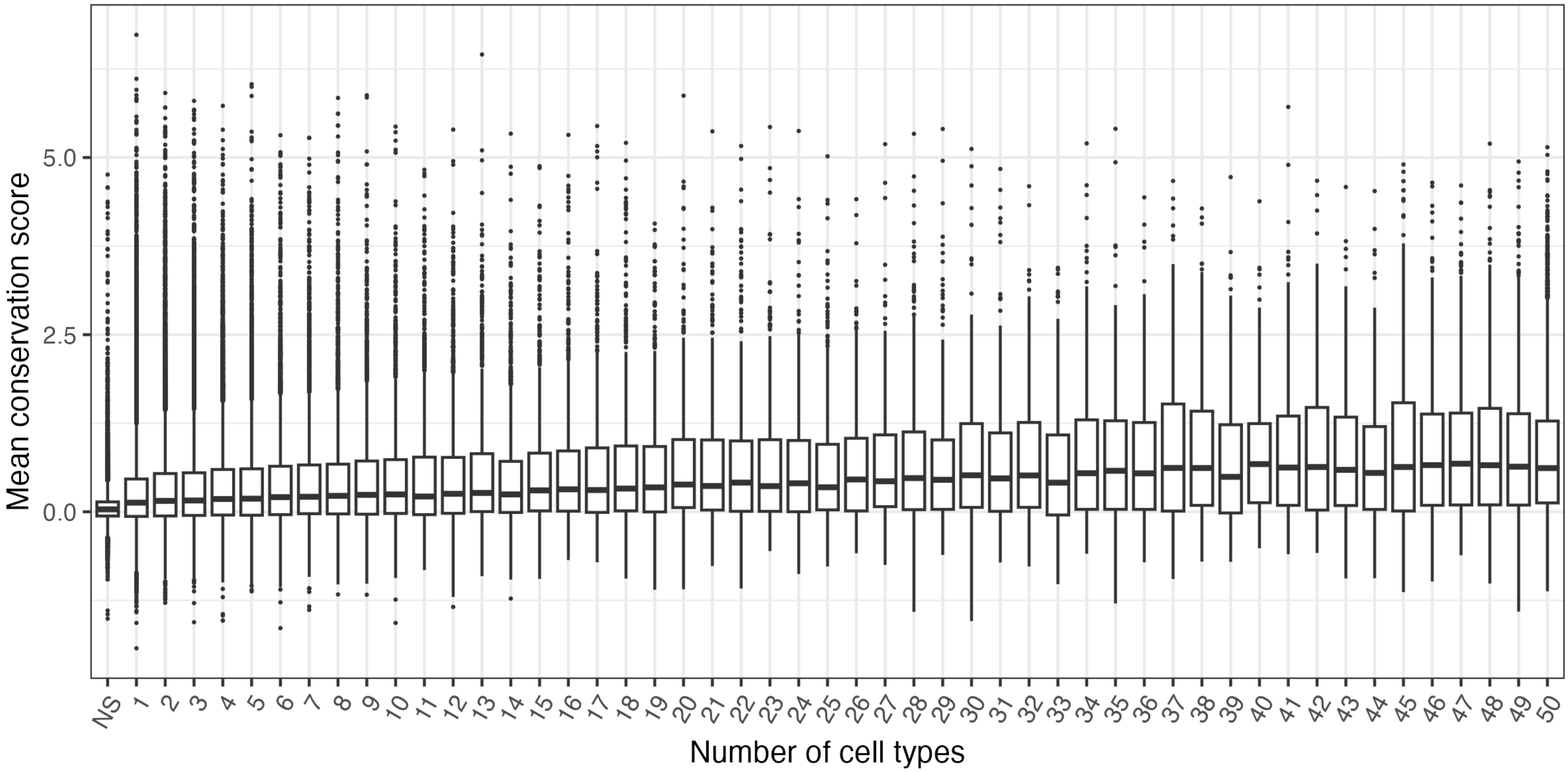


**Supplementary Figure 4.** Conservation scores of CREs across the number of cell types they are active in.

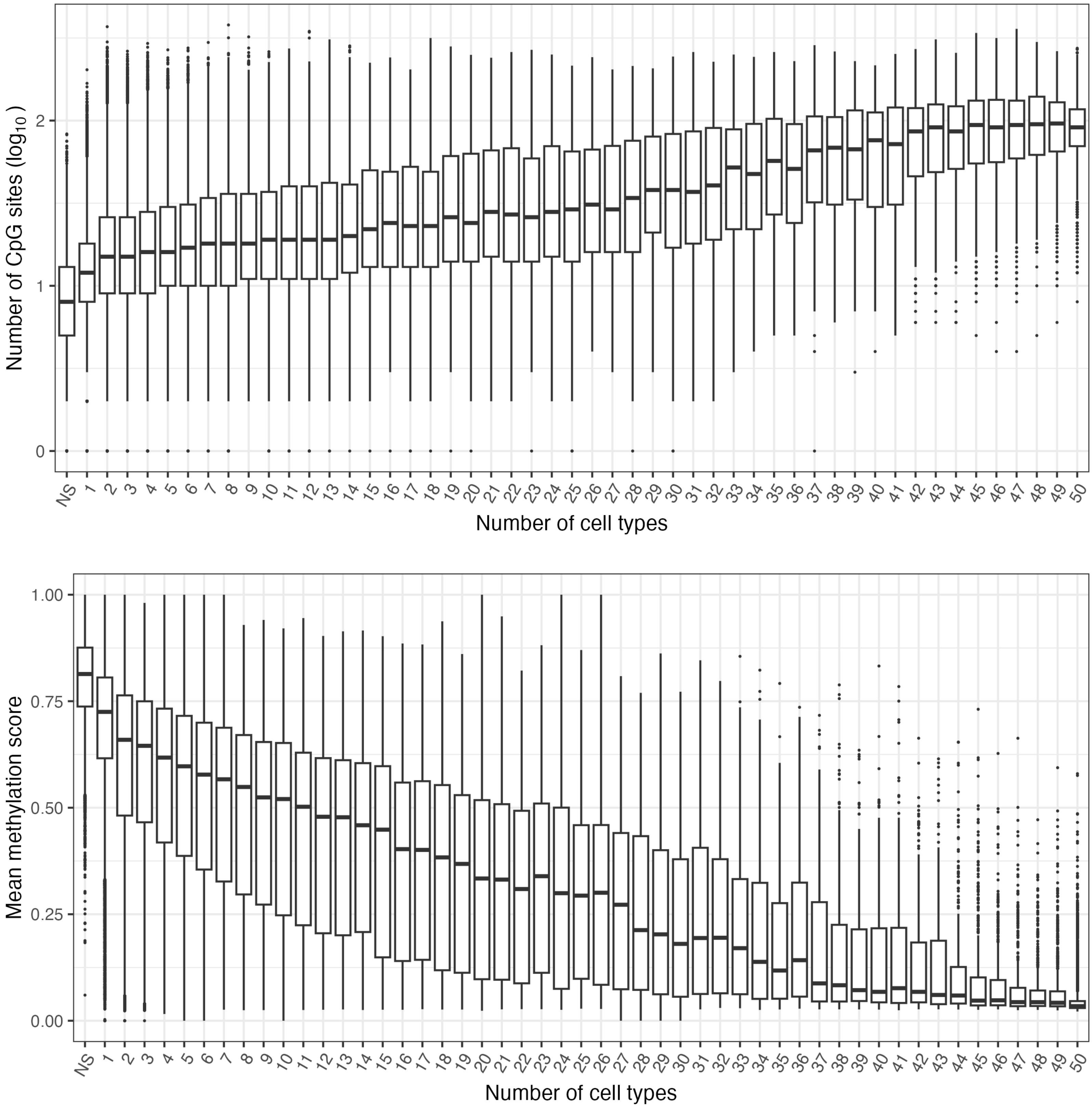


**Supplementary Figure 5.** HK-CREs overlaps with a high number of unmethylated CpG sites as compared to CTS-CREs and negative samples (NS).

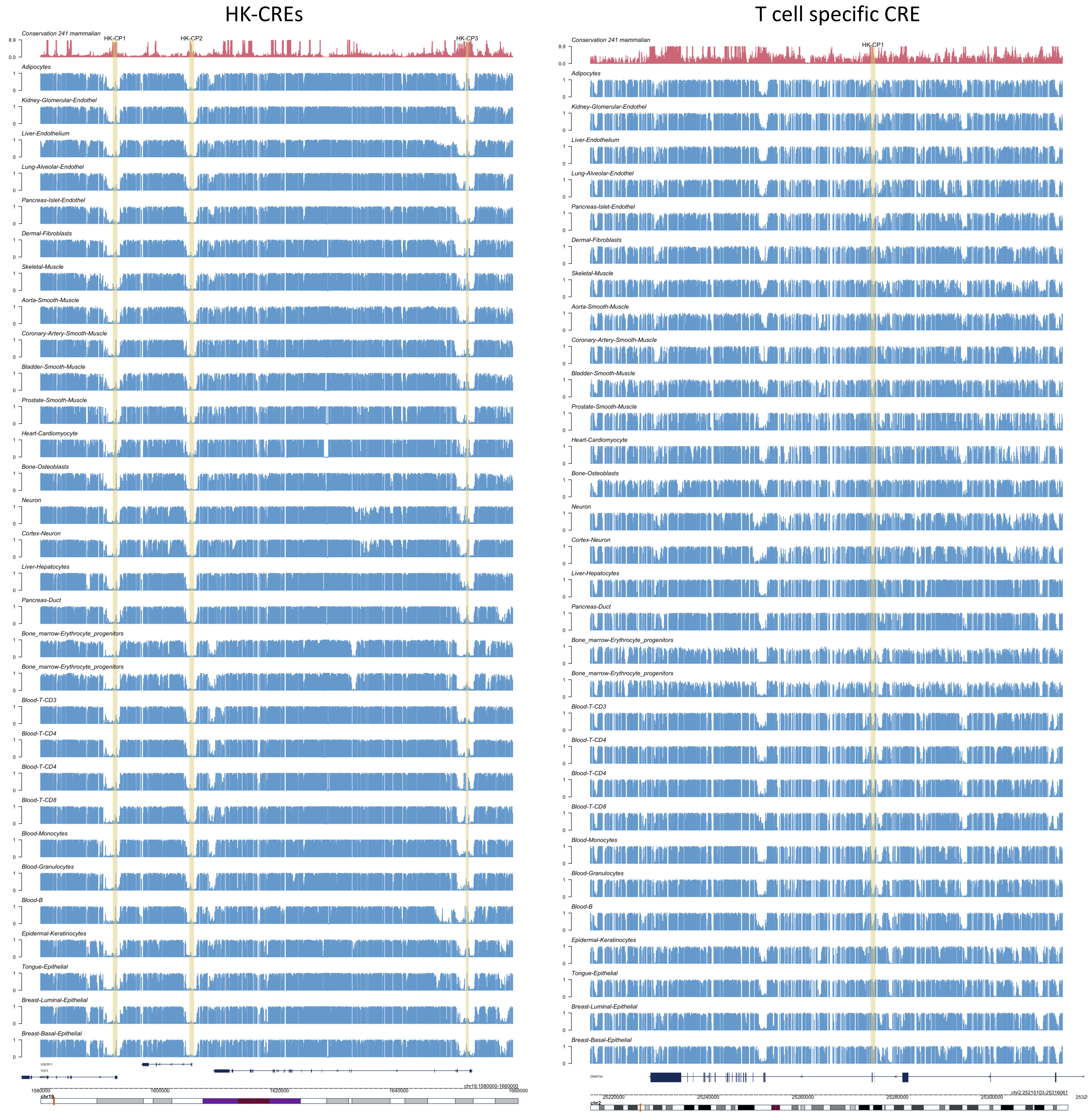


**Supplementary Figure 6.** An example of three HK-CREs shows that they reside within regions dense on CpG sites. The HK-CREs present similar unmethylation patterns across cell types. In contrast, an example of a T cell-specific CREs resides in a region of scarce CpG sites with different methylation patterns across cell types.

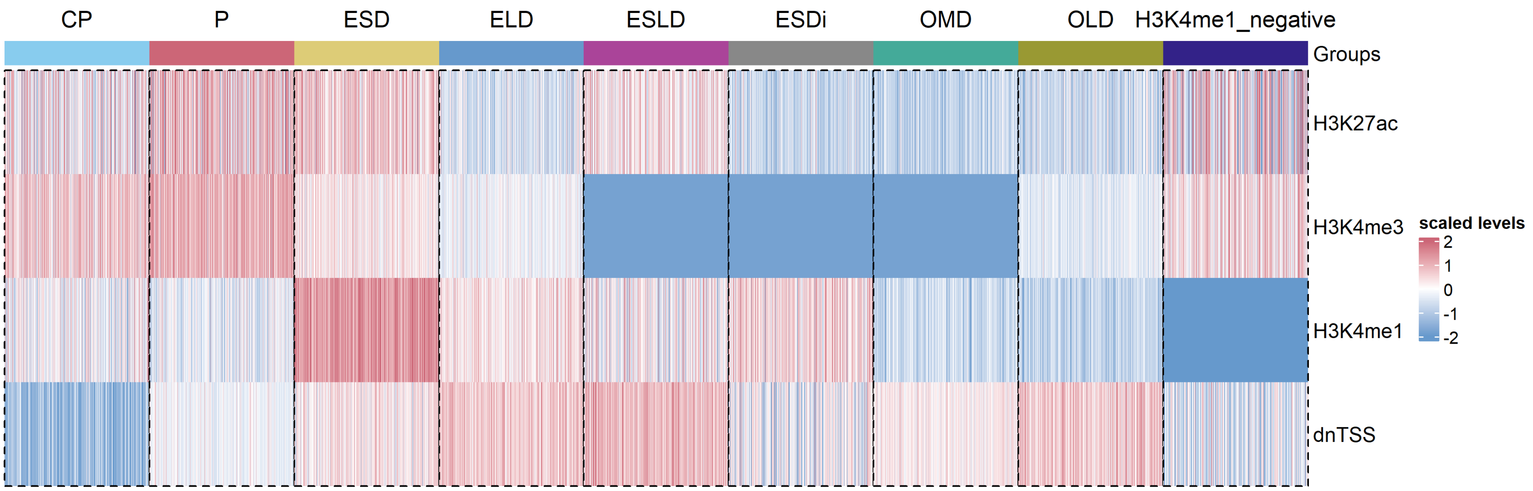


**Supplementary Figure 7.** Heatmap showing the histone modification signals and distance to nearest TSS across clusters.

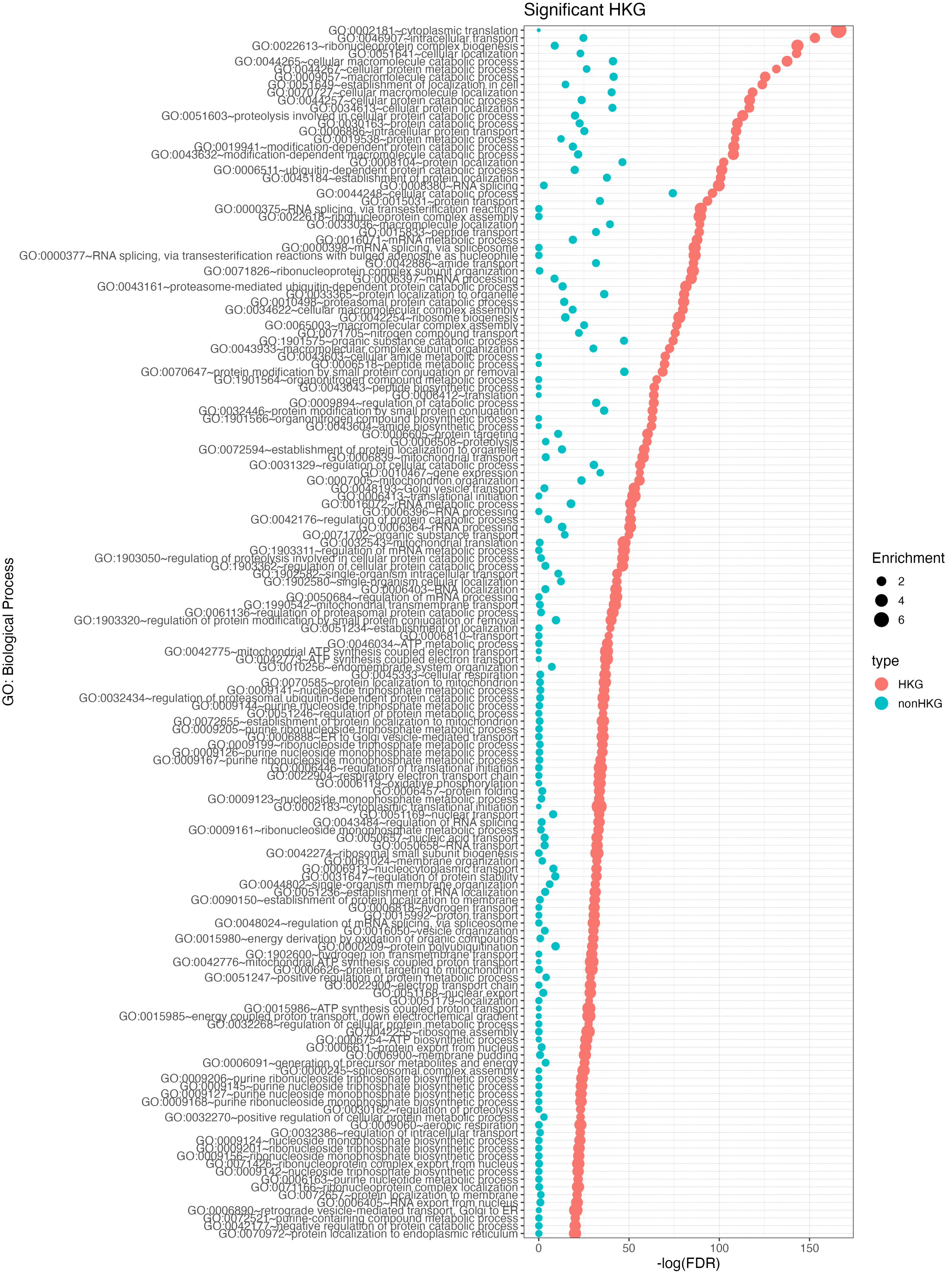


**Supplementary Figure 8.** GO enrichment analysis between HKGs and nonHKGs sorted by significance on HKG. nonHKGs complement HKGs on important housekeeping biological processes.

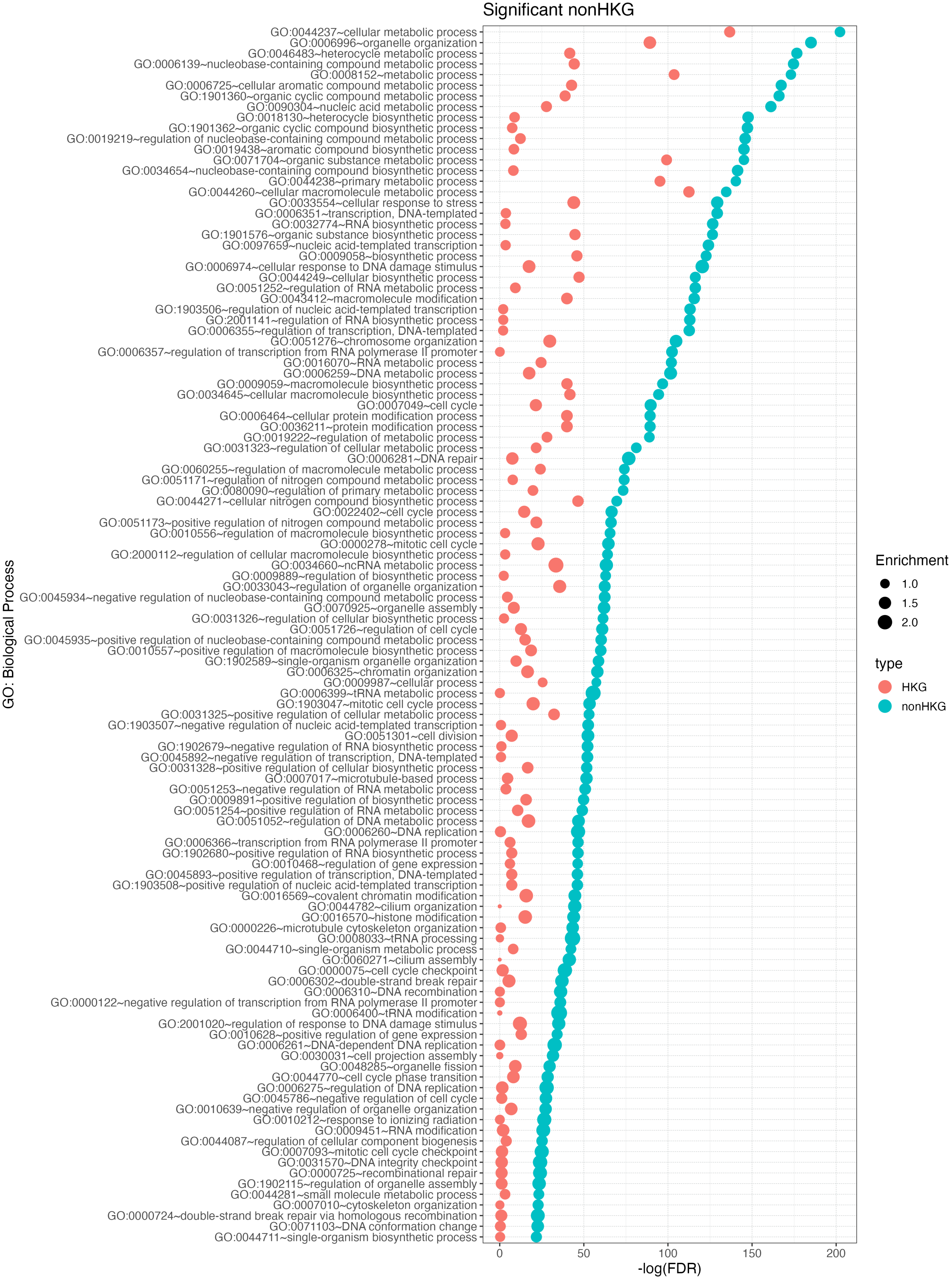


**Supplementary Figure 9.** GO enrichment analysis between HKGs and nonHKGs sorted by significance on nonHKG. nonHKGs complement HKGs on important housekeeping biological processes.

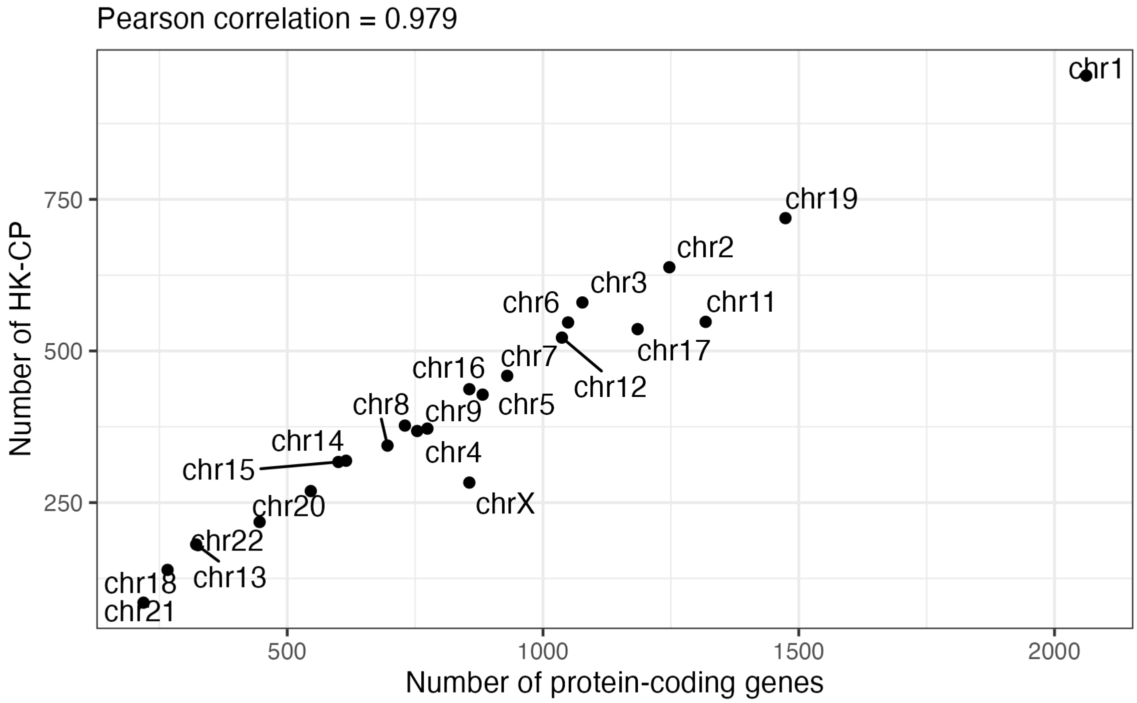


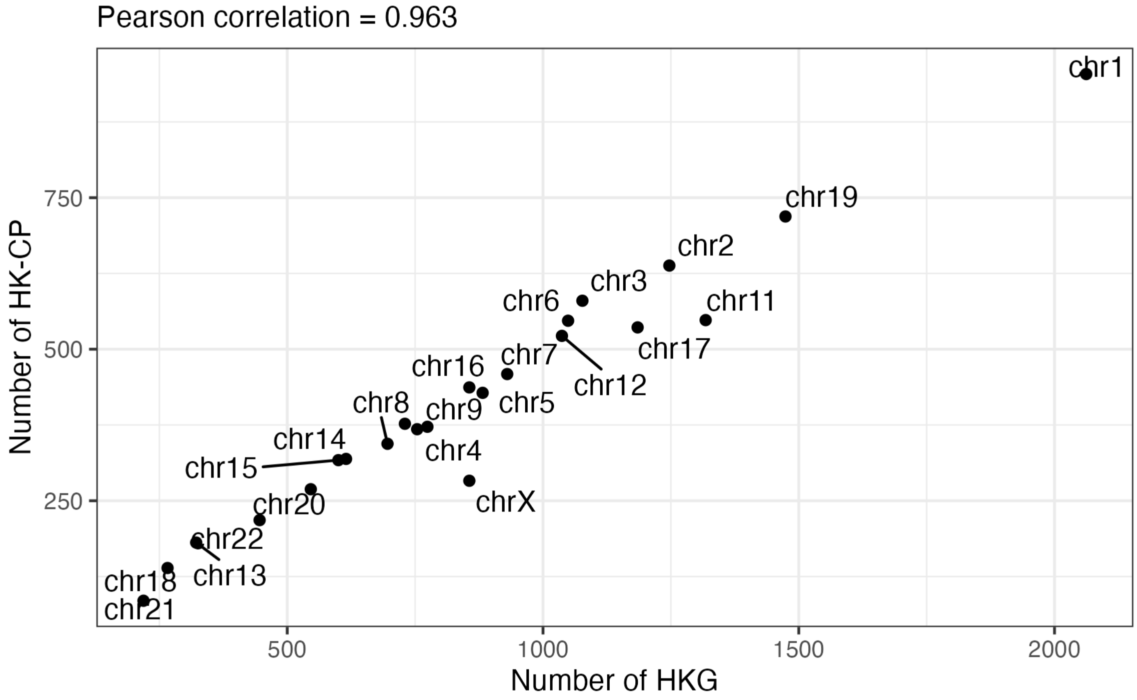


**Supplementary Figure 10.** The number of protein-coding genes and HKG highly correlate with the number of housekeeping core promoters (HK-CP) across chromosomes.

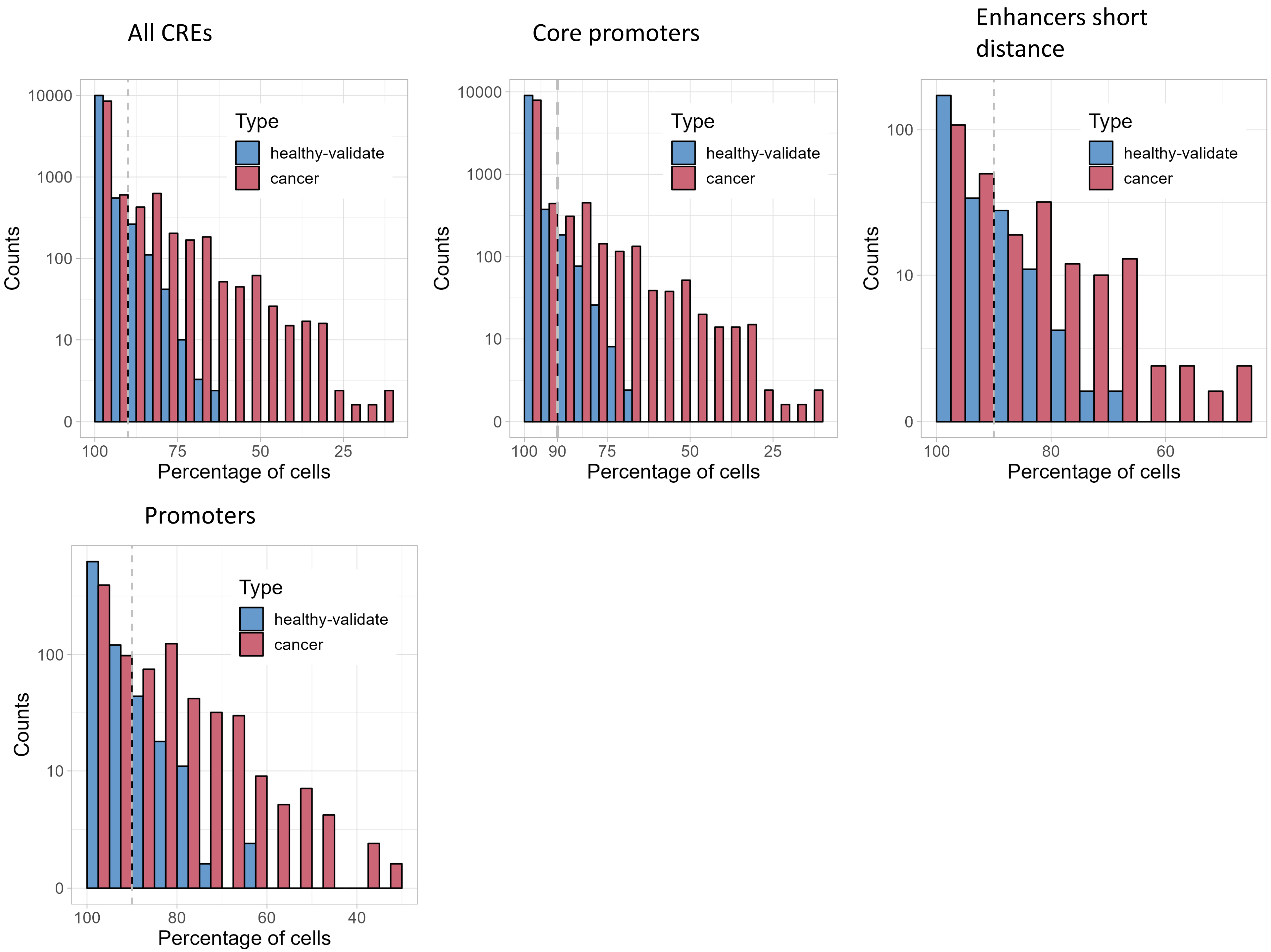


**Supplementary Figure 11.** Validation of housekeeping cis-regulatory elements (HK-CREs) in cancer and healthy cells. The set of HK-CREs was validated in terms of a validation dataset of 21 healthy cell types and a dataset of 26 cancer cell lines. Most of the HK-CREs fulfilled the housekeeping definition in the healthy dataset, e.g. found in at least 90% of the cells. However, the distribution of HK-CREs in cancer cell lines suggests inactive HK-CREs in these cells. The housekeeping core promoters (HK-CPs) were the most affected ones, where a set of HK-CPs were found to be active in less than 35% of the cancer cells.

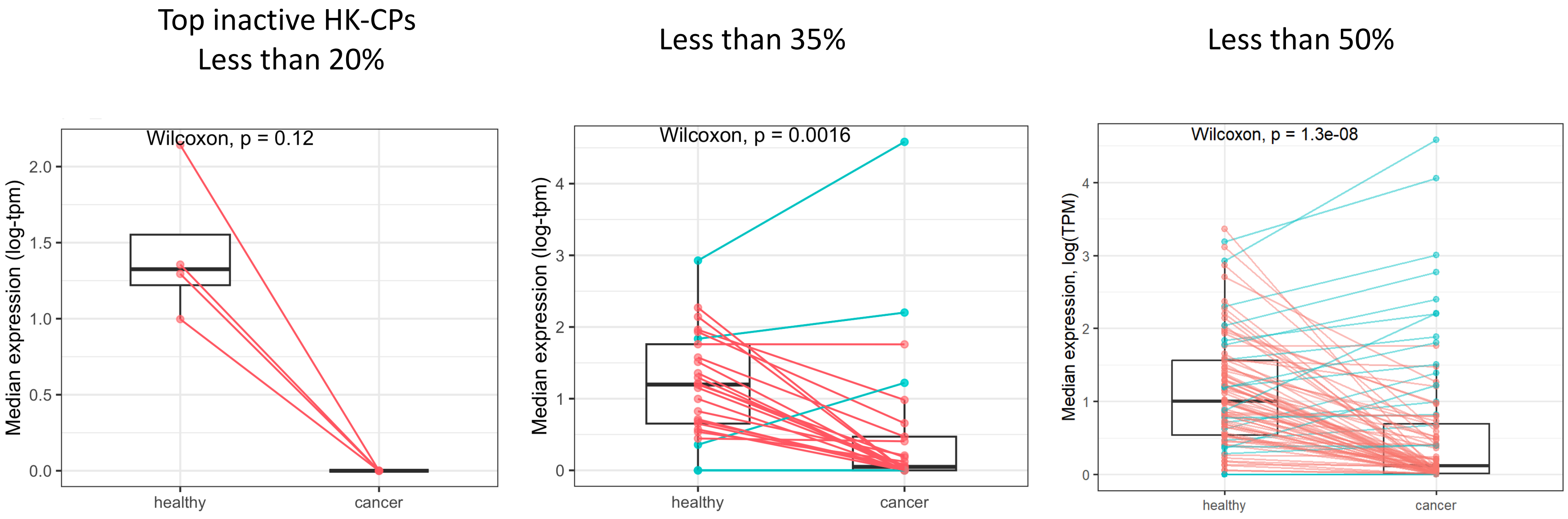


**Supplementary Figure 12.** Comparison of the expression levels of genes whose housekeeping core promoters are inactive in cancer cells. In general, the expression levels are lower in cancer cell lines as compared to healthy cell types. Lines pair the expression levels from healthy to cancer samples and are colored in blue or red, highlighting genes with increased or decreased expression, respectively. The differences are stronger with a stricter selection of genes, e.g. selecting genes active in less than 20%, 35%, or 50%. Genes related to the top inactive HK-CPs (active in less than 20% of cancer cells) present considerably reduced expression levels in cancer cells, where their median expression levels were close to zero.

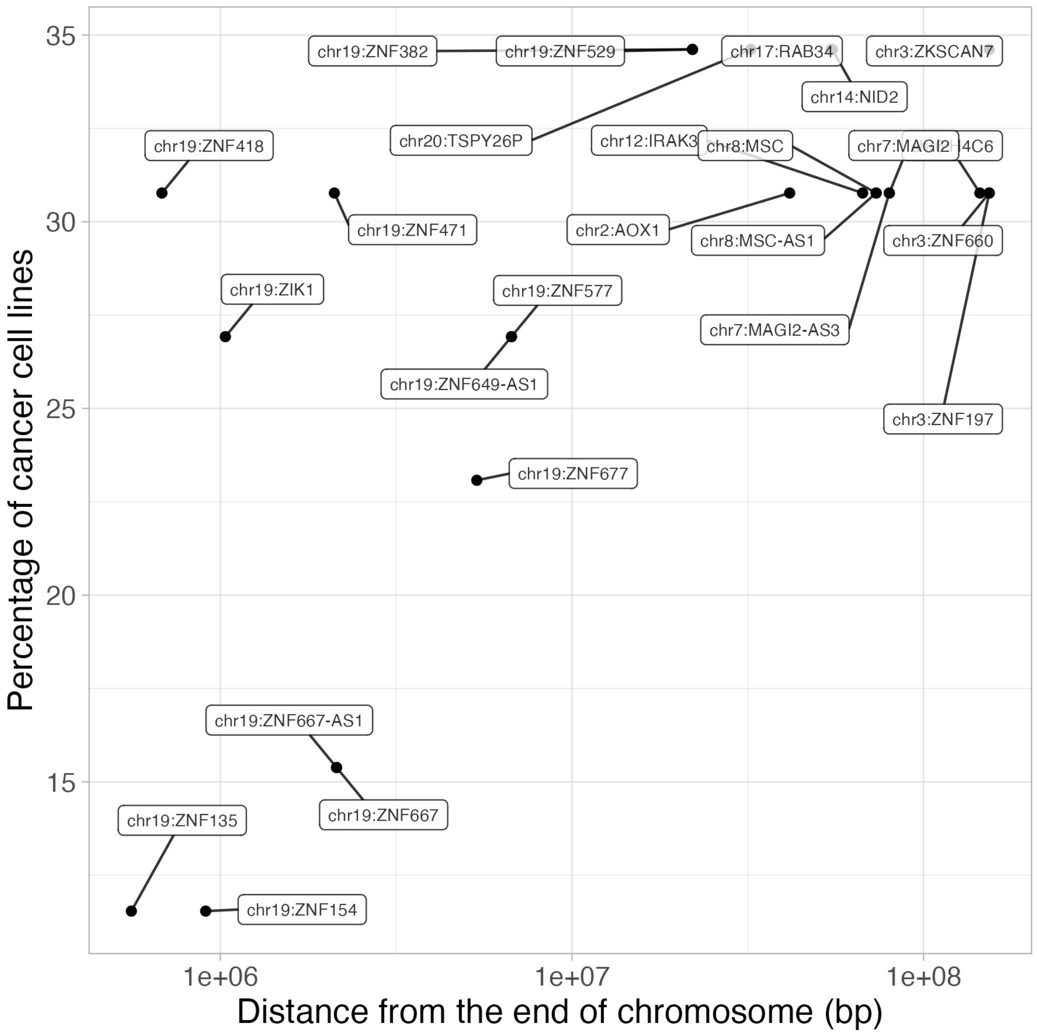


**Supplementary Figure 13.** Comparison of the genes regulated but the top HK-CPs inactive in cancer cell lines by the percentage of cancer cell they are found active and by their distance to the end of the chromosome. Interestingly, chromosome 19 is the most affected one, where genes found active in less than 20% of the cancer cell tend to be located close to the end of the chromosomes, e.g. close to the telomere region, suggesting the importance of this locus in cancer suppression/development.

### ZNF667

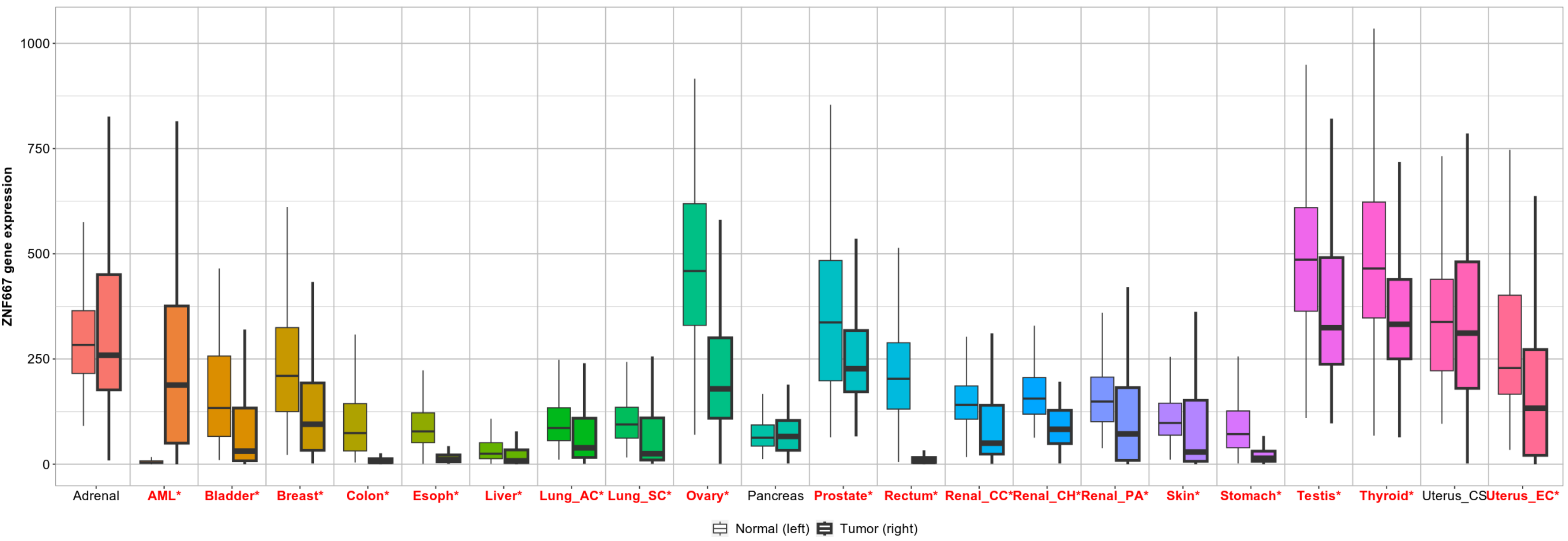


### ZNF667-AS1

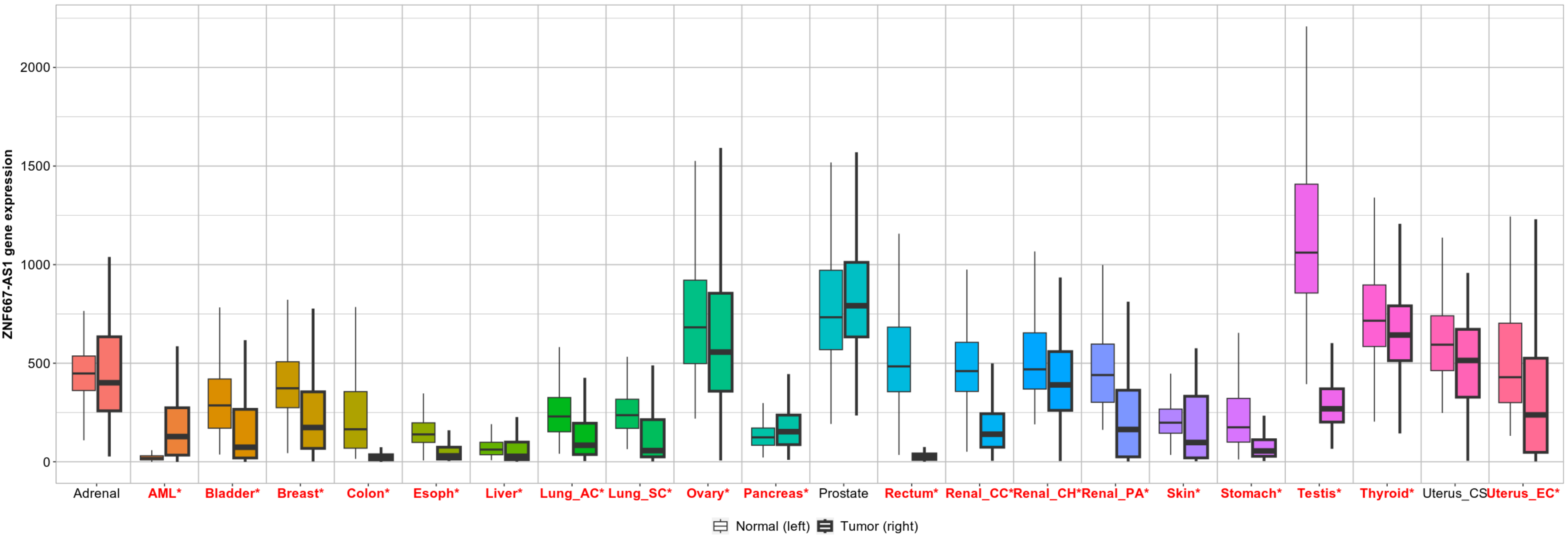


**Supplementary Figure 14.** Comparison of the gene expression levels between tumor and normal samples on different cancer subtypes of the top genes which HK-CP is inactive in cancer cell lines. Red-colored letters indicate a significant difference between tumor and normal samples. In general, the expression levels are lower in tumor samples.

ZNF154

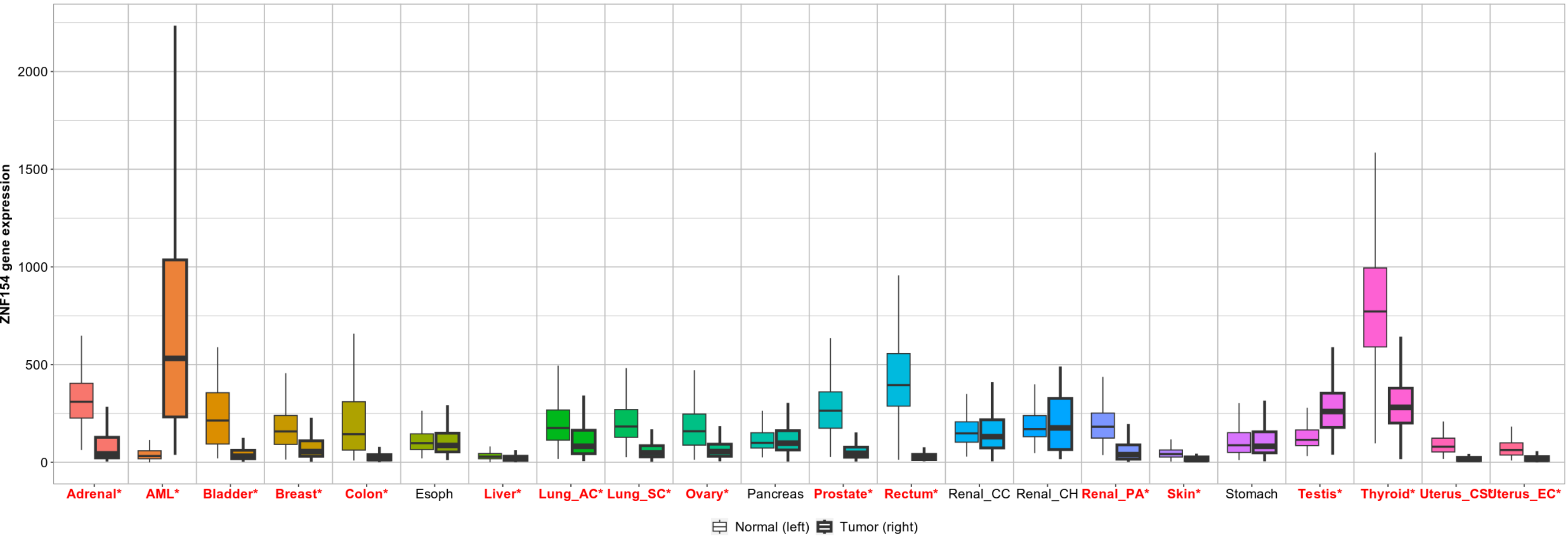


ZNF135

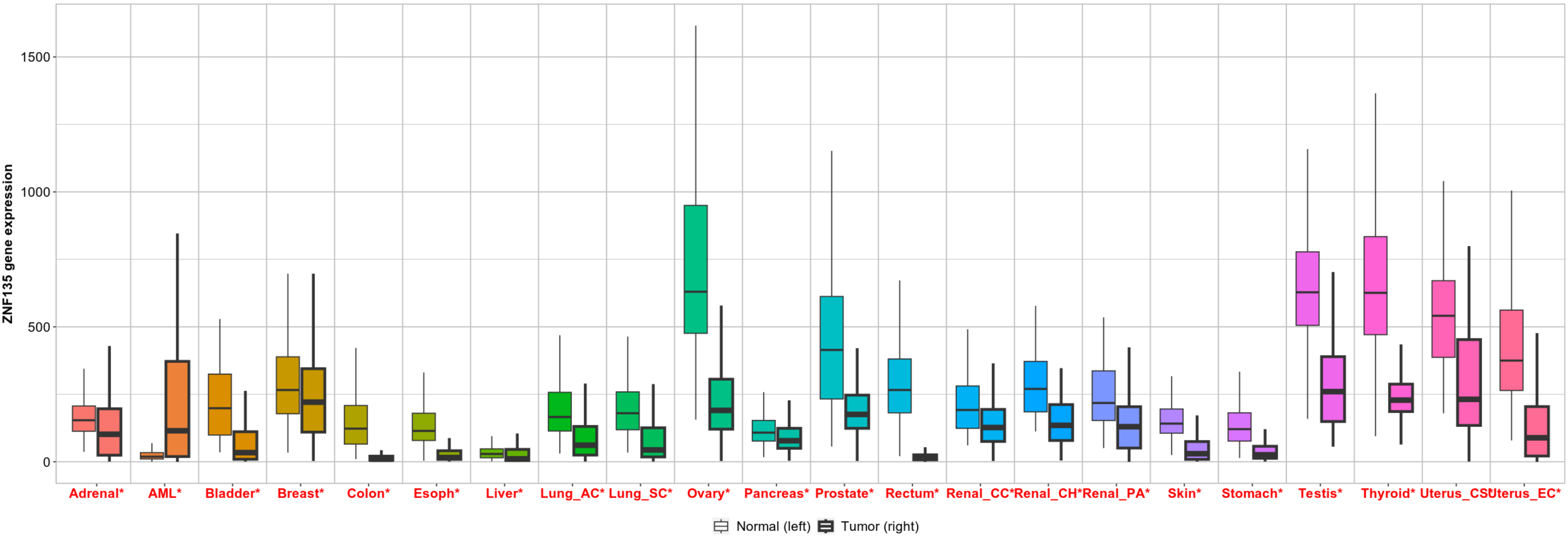


**Supplementary Figure 15.** Comparison of the gene expression levels between tumor and normal samples on different cancer subtypes of the top genes which HK-CP is inactive in cancer cell lines. Red-colored letters indicate a significant difference between tumor and normal samples. In general, the expression levels are lower in tumor samples.

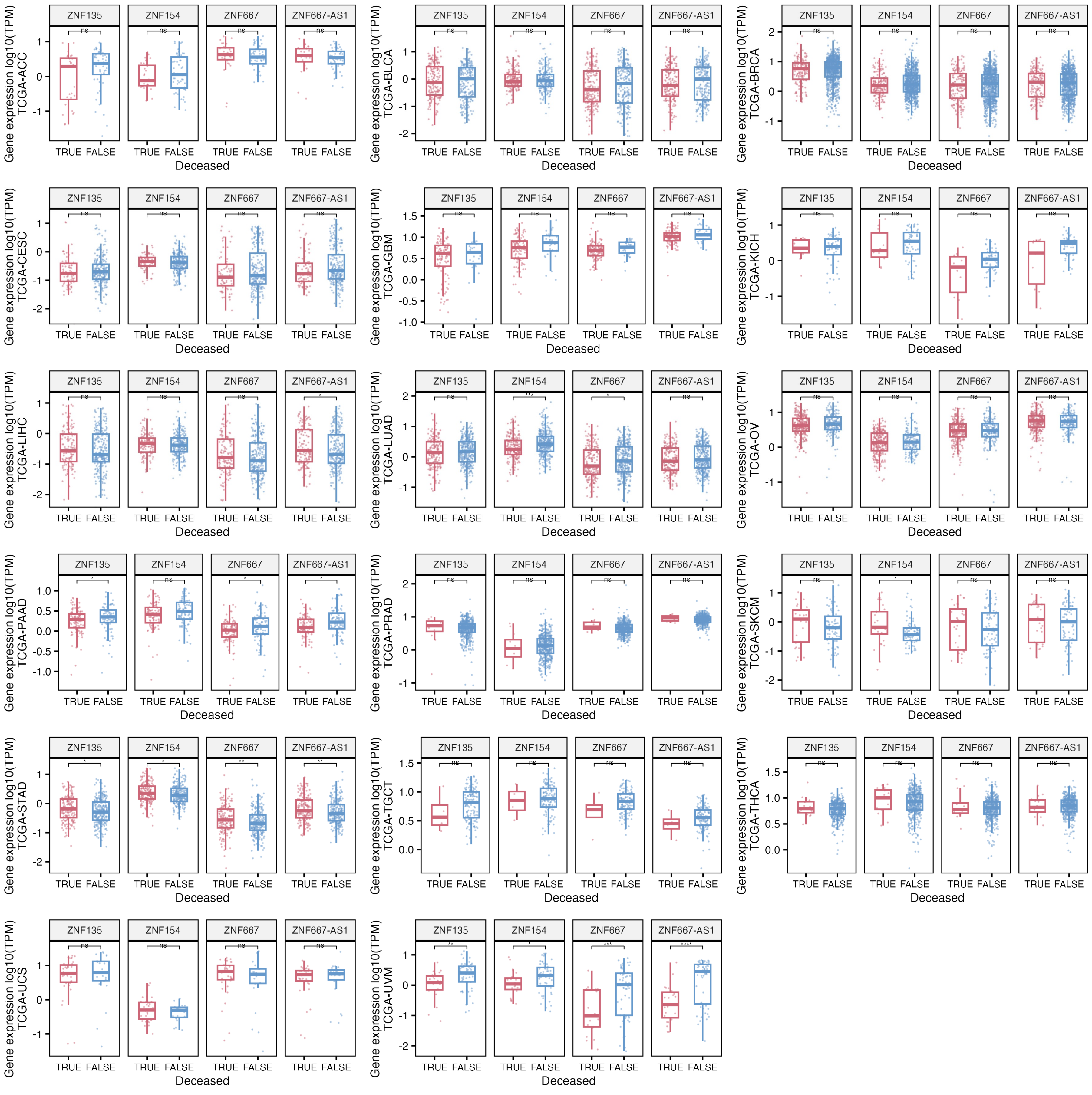


**Supplementary Figure 16.** Comparison of the gene expression of putative tumor suppressor genes: ZNF154, ZNF135, ZNF667, and ZNF667-AS1, by deceased condition across 17 TCGA projects. While most of the comparisons are not significant in all the projects, samples from UVM and PAAD showed a clear upregulation of the genes in non-deceased samples.

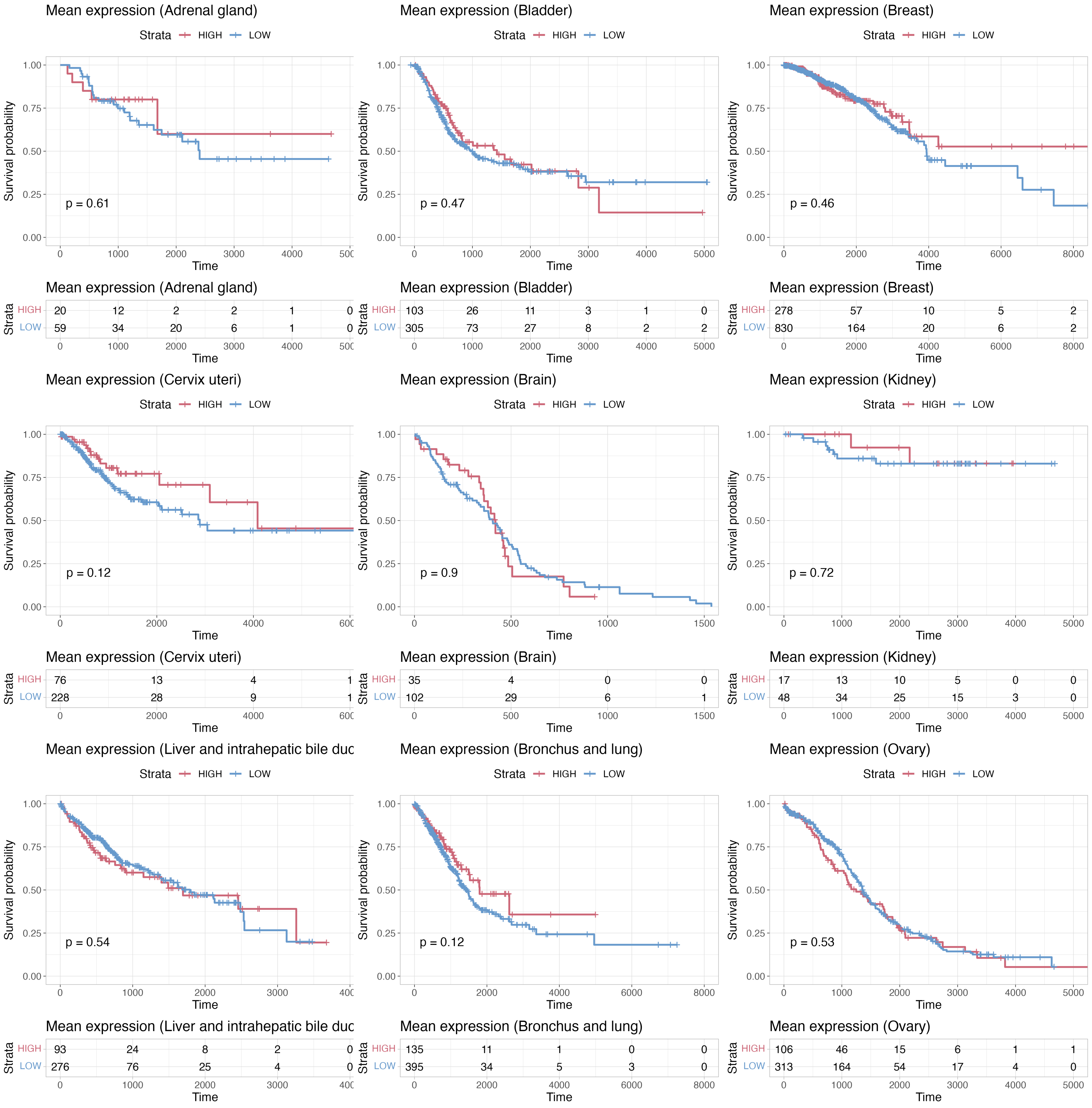


**Supplementary Figure 17.** Independent survival analyses of samples with a high mean expression of putative housekeeping tumor suppressor genes across TCGA projects. While there is an increased survival probability trend of samples with high levels of the genes, the comparisons were not significant.

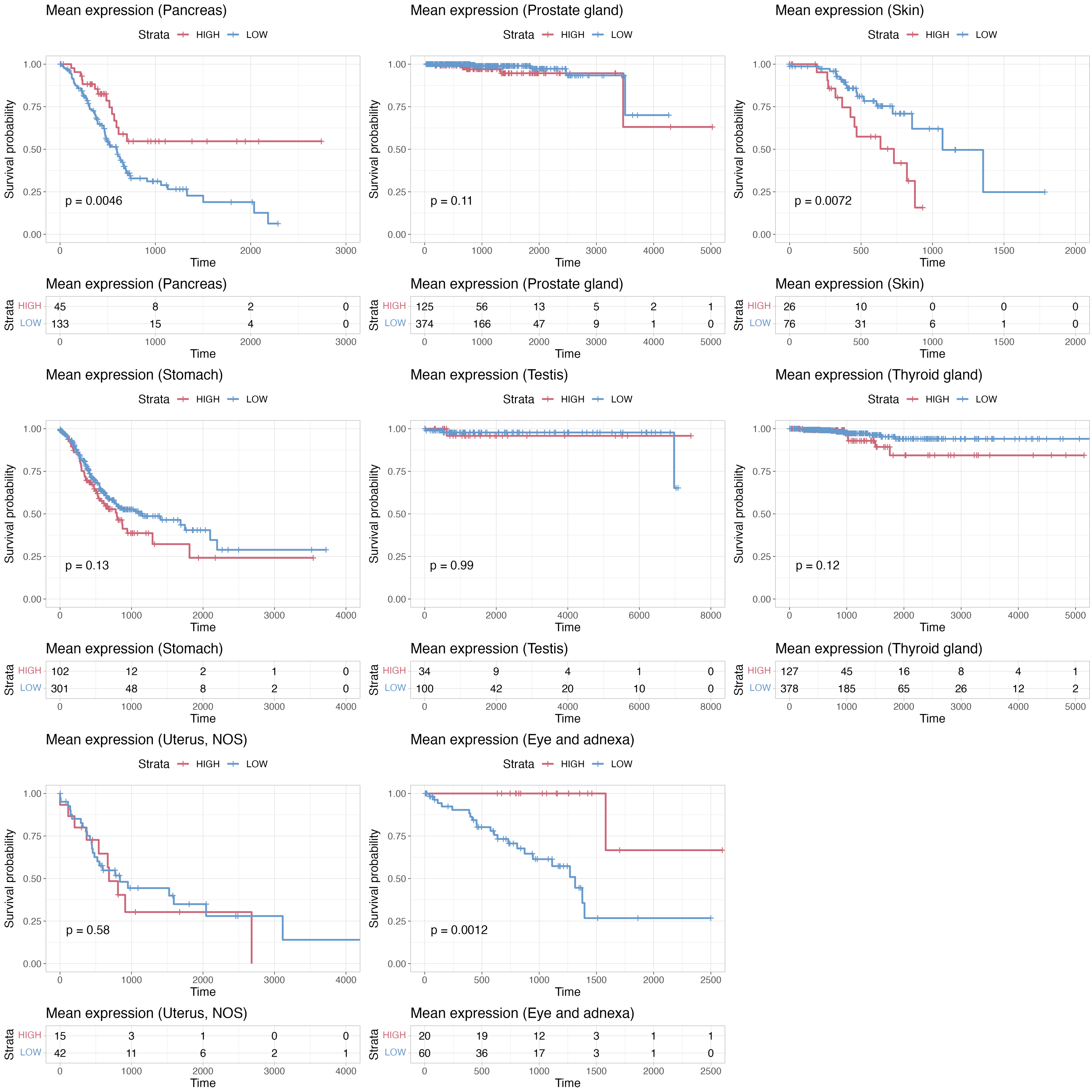


**Supplementary Figure 18.** Independent survival analyses of samples with a high mean expression of putative housekeeping tumor suppressor genes across TCGA projects. Samples from the eye and adnexa, and pancreas samples showed a significant increase in the survival probability of samples with high levels of the genes.

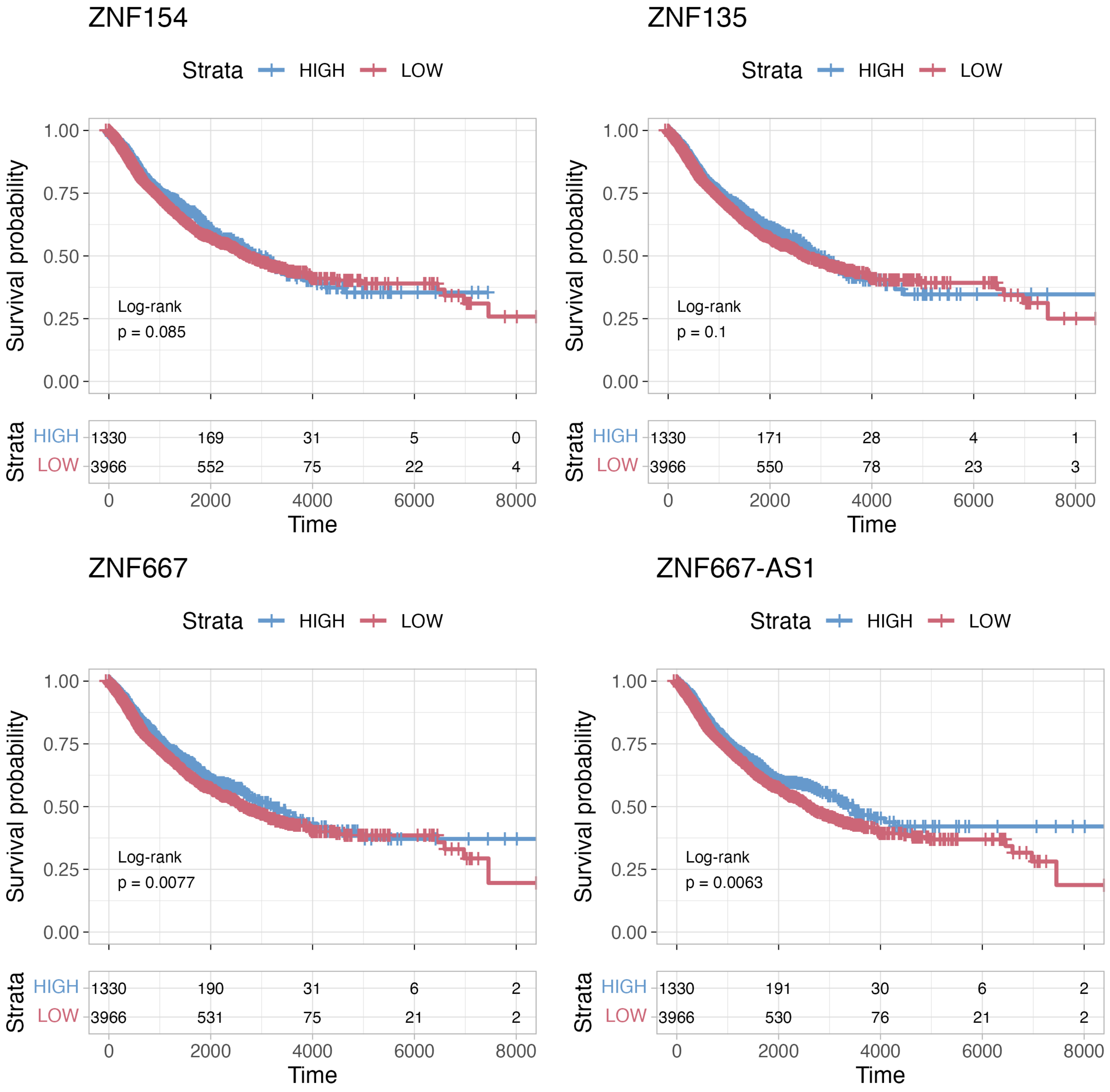


**Supplementary Figure 19.** Independent survival analyses of putative tumor suppressor genes on samples from 17 TCGA projects. All of the analyses showed an increase in the survival probability of samples with high levels of the genes. However, the differences were significant only for ZNF667 and ZNF667-AS1 genes.

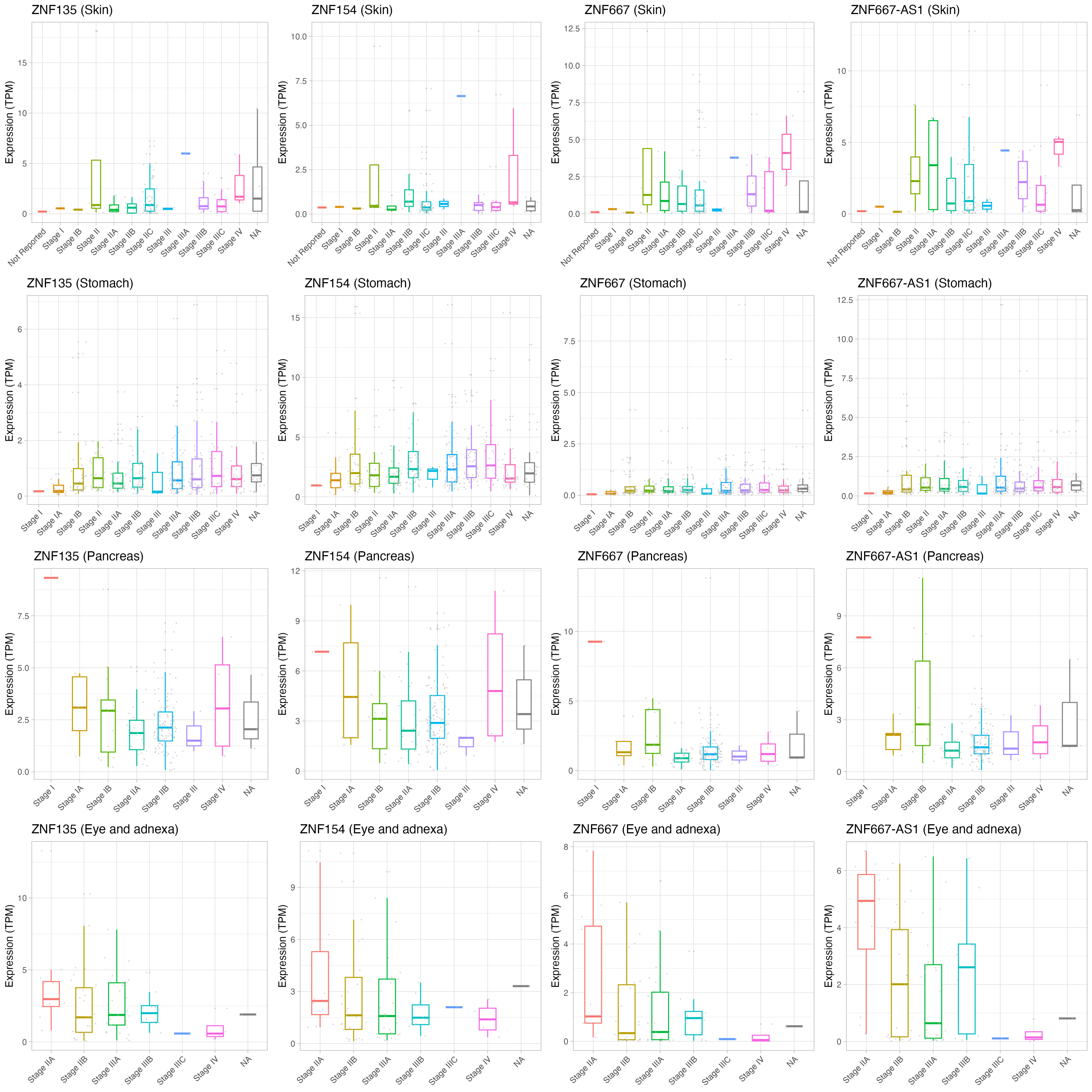


**Supplementary Figure 20.** Comparison of putative tumor suppressor genes across cancer stage in samples from pancreas, eye and adnexa, stomach, and skin TCGA projects. Each project showed a distinct dynamic of the genes, suggesting cancer-specific control of these genes.
